# Targeting the catalytic activity of HDAC1 in T cells protects against experimental autoimmune encephalomyelitis

**DOI:** 10.1101/2023.04.14.536700

**Authors:** Ci Zhu, Valentina Stolz, Natalija Simonovic, Osamah Al-Rubaye, Terezia Vcelkova, Verena Moos, Lena Hess, Astrid Hagelkruys, Moritz Madern, Wolfgang Reiter, Arabella Meixner, Christoph Bock, Markus Hartl, Ellmeier Ellmeier, Christian Seiser

## Abstract

Histone deacetylases are key epigenetic regulators that control T cell-mediated immunity. A T cell-specific deletion of *Hdac1* (HDAC1^cKO^) protects mice against experimental autoimmune encephalomyelitis (EAE). However, it remains elusive whether inhibition of HDAC1 enzymatic activity, which could be achieved therapeutically by HDAC1 inhibitor treatment, is sufficient to block EAE induction. In order to address this question, we generated a novel mouse strain that expresses catalytically inactive HDAC1 (HDAC1^Off^) from the *Rosa26* locus in HDAC1^cKO^ CD4^+^ T cells to mimic selective inhibition of HDAC1 enzymatic activity *in vivo*. Mice expressing wildtype HDAC1 in HDAC1^cKO^ CD4^+^ T cells (HDAC1^On^) were generated as corresponding controls. In contrast to HDAC1^On^ mice, HDAC1^Off^ mice did not develop EAE, and this correlated with diminished leukocyte CNS infiltration. HDAC1^Off^ CD4^+^ T cells in the CNS displayed a severe reduction of IFNγ, IL-17A and TNFα proinflammatory cytokine expression, and *in vivo* activated HDAC1^Off^ CD4^+^ T cells downregulated gene sets associated with T cell activation, cytokine expression and cell migration. This indicates impaired effector functions of HDAC1^Off^ CD4^+^ T cells. Taken together, our study demonstrates that the inhibition of the catalytic activity of HDAC1 in T cells is sufficient to achieve a clinical benefit in EAE disease development. This raises the translational perspective of pharmacological HDAC1 inhibition for treating human T cell-mediated autoimmune diseases.

**Highlights:**

- Successful generation of a novel mouse model that expresses enzymatic-inactive HDAC1 to mimic HDAC1 inhibitor treatment *in vivo*.
- Mice expressing enzymatically inactive HDAC1 instead of WT HDAC1 in T cells do not develop EAE and display diminished leukocyte CNS infiltration.
- *In vivo* activated CD4^+^ T cells expressing enzymatic inactive HDAC1 downregulate pathways important for T cell activation, cytokine expression and cell migration.
- Demonstrate the proof-of-principle that targeting the enzymatic activity of HDAC1 is a promising treatment strategy for autoimmune diseases.

## 1. Introduction

Histone deacetylases (HDACs) are important epigenetic regulators of cell differentiation and homeostasis. They are a large family of enzymes, grouped into four classes, that remove acetyl groups from acetylated lysines of histones as well as non-histone targets [1, 2]. Reversible histone acetylation leads to chromatin remodeling and regulation of gene expression [3, 4]. Reversible acetylation of non-histone targets on the other hand can impact protein stability, localization, function and protein-protein interactions [5]. Class I HDACs comprise four enzymes, namely HDAC1, HDAC2, HDAC3 and HDAC8 [6, 7]. HDAC1 and HDAC2, together or alone, form the catalytic core of multi-protein co-repressor complexes including SIN3, NURD and COREST [8–11].

HDAC inhibitors are currently used as anti-cancer agents [12–15]. Additional studies using preclinical mouse autoimmune models indicated that HDAC inhibitors are promising therapeutic drugs to treat T cell-mediated diseases [16–18]. Experimental autoimmune encephalomyelitis (EAE) is an animal disease model for autoimmune central nervous system (CNS) inflammation and resembles in certain aspects multiple sclerosis (MS) in humans [19, 20]. Treatment of mice with pan-HDAC inhibitors such as trichostatin A [21, 22] and vorinostat [23] leads to an amelioration of EAE. However, due to the broad action of pan-HDAC inhibitors, these drugs can lead to severe side effects and their clinical use is therefore limited [24]. Specific inhibitors of class I HDACs have shown therapeutic potential in preclinical trials although no human clinical data are available yet for inflammatory and autoimmune diseases [25]. We have recently shown that conditional knockout mice lacking HDAC1 in T cells (using the *Cd4*-Cre deleter strain; *Hdac1*^fl/fl^ x *Cd4*-Cre; HDAC1^cKO^) are protected from EAE disease development [26]. We further revealed that HDAC1 is an important regulator for T cell trafficking in EAE, since HDAC1-deficient CD4^+^ T cells fail to migrate into the CNS [27]. These results suggest that HDAC1 might be a suitable target for treating autoimmune CNS diseases including MS.

In order to further test the translational potential of our finding that HDAC1 is a key target to prevent EAE development in a more clinical situation, we generated a novel mouse strain that allows to conditionally replace the expression of endogenous HDAC1 wildtype (WT) enzyme by the *Rosa26* locus-driven expression of a transgenic, catalytically inactive version of HDAC1 in T cells. Using these mice, we demonstrated that mice expressing catalytically inactive HDAC1 are protected from EAE disease development. In contrast, control mice expressing a transgenic WT HDAC1 version from the *Rosa26* locus were susceptible to EAE induction. While catalytically inactive HDAC1 and wildtype HDAC1 were equally well incorporated into co-repressor complexes and showed very similar interactomes, RNA sequencing revealed that inactivation of HDAC1 catalytic activity led to a decreased inflammatory phenotype of CD4^+^ T cells *in vivo*. Further, the inactivation of HDAC1 catalytic activity resulted in the downregulation of pathways important for T cell activation and migration. In summary, by using our novel mouse models we showed that the catalytic activity of HDAC1 is a promising target for the treatment of EAE.

## 2. Material and Methods

### 2.1. Animal models

Animal experiments were evaluated by the ethics committee of the Medical University of Vienna and approved by the Austrian Federal Ministry for Education, Science and Research (animal protocol numbers 66.009/0039-V/3b/2019, 66.009/0041-V/3b/2019, 66.009/0326-V/3b/2019). Animals were maintained at the Medical University of Vienna core facility laboratory animal breeding and husbandry and in research facilities of the Max Perutz Labs at the Vienna BioCenter. Animal husbandry and experiments were performed under national laws in agreement with guidelines of the Federation of European Laboratory Animal Science Associations (FELASA), which correspond to Animal Research: Reporting of *in vivo* Experiments from the National Center for the Replacement, Refinement and Reduction of Animals in Research (ARRIVE) guidelines. *Cd4*-*Cre* deleter mice [28] (MGI:2386448) were kindly provided by Christopher Wilson (University of Washington, Seattle). 2D2 TCR transgenic mice (2D2) recognizing MOG_35−55_ peptide [29] (MGI:3700794) were kindly provided by Gernot Schabbauer (MedUni Vienna, Austria). CD45.1^+^ and *Rag2*^−/−^ mice were kindly provided by Jochen Huehn (Helmholtz Center, Braunschweig, Germany). *Hdac1*^fl/fl^ x *Cd4*-Cre mice (HDAC1^cKO^) have been previously generated [30]. All mice analyzed were 8-12 weeks of age and of mixed sex, unless stated otherwise.

### 2.2. Generation of transgenic *Rosa26* mice

Cloning of constructs for targeting the murine *Rosa26* locus: For expression of FLAG-tagged versions of wildtype HDAC1 (i.e. HDAC1^On^) and catalytically-inactive HDAC1 (i.e. HDAC1^Off^), the open reading frame (ORF) of mouse HDAC1 was cloned into the FLAG-tag expression vector pCMV-Tag4 and the codon for H141 was mutagenized to the alanine codon GCC. Expression of HDAC1^On^ and HDAC1^Off^ proteins was confirmed upon transient transfection in HeLa cells and stable expression in HAP1 cells [31]. The ORFs encoding FLAG-tagged HDAC1 WT and HDAC1 CI were cloned into the exchange vector pBTV2-11. The resulting exchange vectors contain a βGEO-STOP cassette and the ORF for FLAG-tagged HDAC1 isoforms flanked by two attB integrase attachment sites [32] (see Fig. 1A for a schematic map of the targeting scheme).

**Figure 1.**
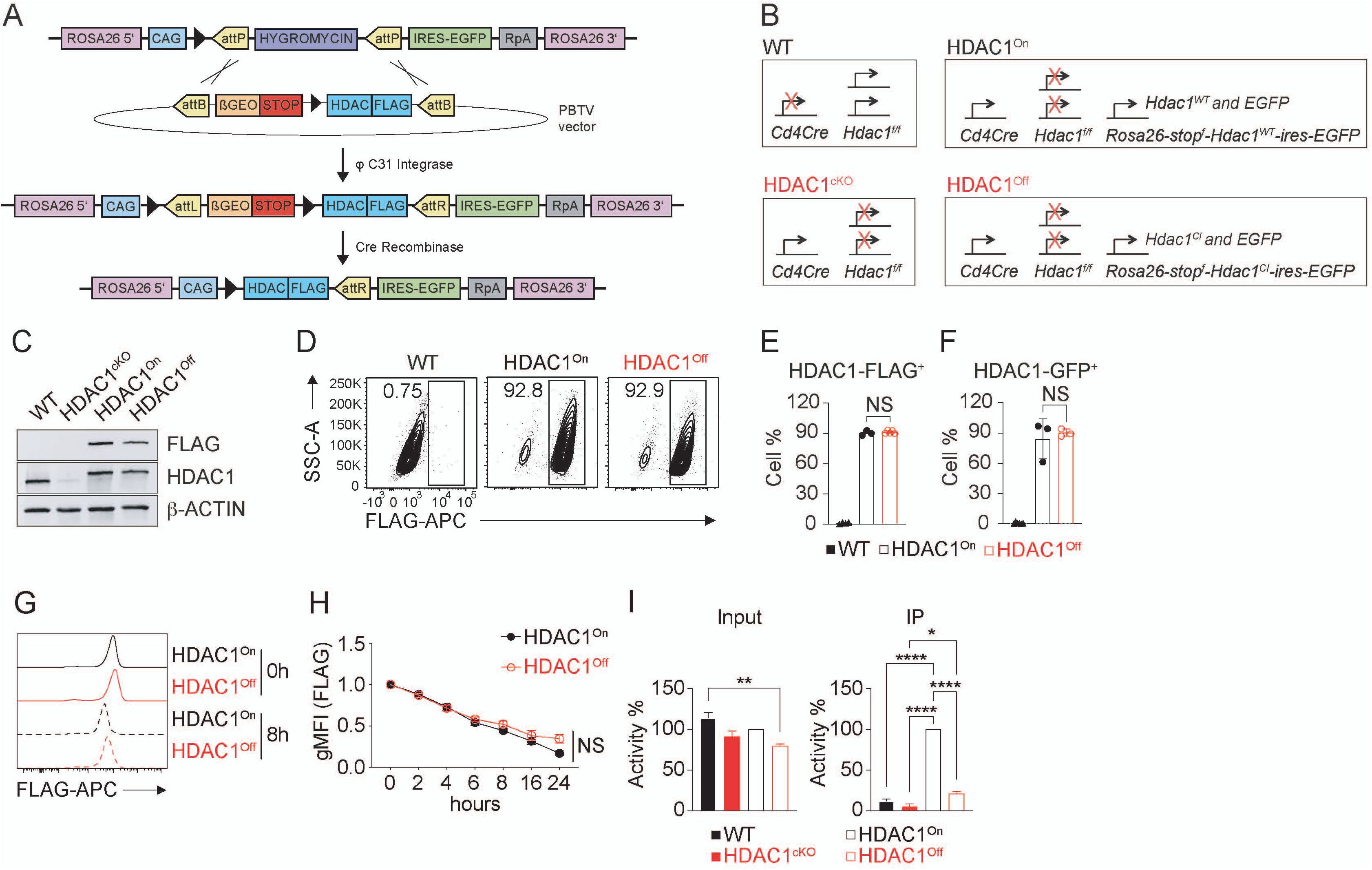
Setup of a conditional mouse model for T-cell specific inactivation of HDAC1 catalytic activity. (**A**) Schematic showing the generation of transgenic mice by ΦC31 integrase-mediated insertion of HDAC1 open reading frames with a C-terminal FLAG tag followed by an IRES-EGFP reporter from the PBTV shuttle vector into the targeting platform at the murine *Rosa26* locus. Deletion of the loxP-flanked STOP cassette by *Cd4*-*Cre* results in the expression of transgenic FLAG-tagged HDAC1 isoforms in T cells. LoxP sites are shown as black triangles. (**B**) Genotypes of mice used in this study: WT (*Cd4-Cre^–^*),HDAC1^On^ (*Hdac1^fl/fl^ Rosa26-STOP^fl^-HDAC1^WT^ Cd4-Cre^+^*) or HDAC1^Off^ (*HDAC1^fl/fl^ Rosa26-STOP^fl^-HDAC1^CI^ Cd4-Cre^+^*). **(C)** Western blot analysis of protein extracts isolated from WT, HDAC1^cKO^, HDAC1^On^ and HDAC1^Off^ Th1 cells using antibodies specific for FLAG, HDAC1 and β-ACTIN. **(D)** Flow cytometry analysis showing the expression of FLAG in WT (= negative control), HDAC1^On^ and HDAC1^Off^ Th1 cells. Numbers indicate the percentages of cells in the respective quadrants. **(E)** Summary showing percentage of FLAG-epitope^+^ cells as assessed in (D). Mean ± SEM is shown. N.S. not significant. Each symbol indicates one mouse. **(F)** Summary showing the frequency of EGFP^+^ WT, HDAC1^On^ and HDAC1^Off^ Th1 cells. (**G**) Flow cytometry analysis showing the steady-state levels of FLAG-tagged HDAC1^On^ and HDAC1^Off^ proteins in activated CD4^+^ T cells after 0 and 8 hours of cycloheximide treatment. Expression was detected using an anti-FLAG antibody. (**H**) Summary showing the expression of FLAG-tagged HDAC1^On^ and HDAC1^Off^ proteins in activated CD4^+^ T cells in the course of 24 hours of cycloheximide treatment. (**I**) Analysis of HDAC1 catalytic activity towards acetylated histones in total cell extracts (Input, left graph) and in FLAG immunoprecipitates (IP, right graph) in WT, HDAC1^cKO^, HDAC1^On^ and HDAC1^Off^ Th1 cells. Mean ± SEM is shown. *p < 0.05, **p < 0.01, and ***p < 0.001 (ordinary 1-way ANOVA analysis followed by Tukey’s multiple-comparisons test). Data are representative (C,D,G) or show the summary (E,F,H) of at least 3 mice or of (I) 3 independent experiments.

ESC transfection and blastocyst injection: The murine embryonic stem cell (ESC) line A9 with the integrated docking platform at the *Rosa26* locus has been generated by homologous recombination with the docking vector containing 5’ and 3’ *Rosa26* homologous arms of the genomic *Rosa26* locus. The docking platform contains the CAG (CMV early enhancer/chicken β-actin) promoter, the 5’ loxP site, the hygromycin resistance gene flanked by two attP integrase attachment sites [32] and the reporter gene IRES-EGFP-RpA. A9 ESCs harboring the docking platform at the *Rosa26* locus were transfected by nucleofection with integrase φC31 plasmid and the respective exchange plasmid using an Amaxa 4D Nucleofector (*Lonza*). The integration leads to replacement of the hygromycin resistance gene by the βGEO-STOP cassette and the HDAC1-FLAG ORF. The STOP signal prevents the expression of HDAC1 due to premature transcriptional termination. Successfully targeted ESCs acquire G418 resistance, mediated by the β-geo gene, which allows the selection of positive clones. After 24 hours of recovery, cells were selected with G418 sulphate at a final concentration of 300 μg/ml for 10-14 days. Successfully targeted clones were identified by PCR and Southern blot analysis and correctly targeted A9 ESCs were injected into C57BL6 blastocysts (IMBA, transgenic service department) as described previously [33]. Transgenic animals were identified by PCR genotyping.

Generation of HDAC1^On^ and HDAC1^Off^ mice: By intercrossing the transgenic *Rosa26* knock-in mouse models with the previously described T cell-specific conditional HDAC1 knockout strain (*Hdac1*^f/f^ *Cd4*-Cre) [30] we generated HDAC1^On^ (*Hdac1^fl/fl^ Rosa26-STOP^fl^-HDAC1^WT^ Cd4-Cre^+^*) or HDAC1^Off^ (*HDAC1^fl/fl^ Cd4*-Cre^+^ *Rosa26-STOP^fl^-HDAC1^Off^ Cd4-Cre^+^*) mice (Fig. 1B).

Primers used for cloning and mutagenesis (sequence 5’ to 3’): BamHI_mHD1_F: GGC GGA TCC ATG GCG CAG ACT CAG GGC A; EcoRI_mHD1_R: ATC GAA TTC GGC CAA CTT GAC CTC TTC T; HDAC1_MUT_F: ACT GGG CCG GGG GCC TGC ACG CCG CAA AGA AGT CTG AAG CTT C; HDAC1_MUT_R: GAA GCT TCA GAC TTC TTT GCG GCG TGC AGG CCC CCG GCC CAG T; XbaI_3’FLAG_R: CCA TCT AGA TTG GGT ACA CTT ACC TGG TAC;

### 2.3. Genotyping of mice

The PCR for the insertion of the *Hdac1* transgene (both catalytically inactive and WT) (KI band: 346bp. WT band: 246bp) was carried out for 5 min at 95°C, followed by 40 cycles of 30 sec at 95°C, 30 sec at 56°C, and 2 min at 72°C. The PCR for the floxed *Hdac1* gene (floxed band: 346bp. WT band: 246bp) was carried out for 5 min at 95°C, followed by 40 cycles of 30 sec at 95°C, 30 sec at 54°C, and 100 sec at 72°C. The PCR for the *Cd4*-Cre transgene (300bp) was carried out for 5 min at 96°C, followed by 39 cycles of 30 s at 94°C, 30 s at 56°C, and 1 min at 72°C. The PCR for the Rosa26 locus (450 bp) was carried out for 5 min at 95°C, followed by 35 cycles of 30 sec at 95°C, 30 sec at 56°C, and 80 sec at 72°C. The PCR for the *2D2* transgene (346 bp) was carried out for 1 min at 94°C, followed by 34 cycles of 30 s at 94°C, 30 s at 60°C, and 45 s at 72°C.

Primers used for genotyping (sequence 5’ → 3’): Hdac1 transgene F: GCA TCG CCT TCT ATC GCC TTC; Hdac1 transgene R: CTT GGT CAT CTC CTC AGC ATT GG; Hdac1 floxed F: GGT AGT TCA CAG CAT AGT ACT T; Hdac1 floxed R: CCT GTG TCA TTA GAA TCT ACT T; Cd4-Cre F: TCT CTG TGG CTG GCA GTT TCT CCA; Cd4-Cre R: TCA AGG CCA GAC TAG GCT GCC TAT; Rosa26 locus F: AAG AAC TGC AGT GTT GAG GC; Rosa26 locus R: TCT CCC AAA GTC GCT CTG AG; 2D2 transgene F: AGG ATG TGA GGG ACT ACC TCC TGT A; 2D2 transgene R: TCC TTC ACT CTG ATT CTG GCA ATT T;

### 2.4. Purification of CD4^+^ T Cells

Cells from spleen, axillary, brachial, and inguinal lymph nodes were isolated and pooled. Single cell suspensions were made using a 70μm cell strainer (Corning) in a 6-well plate (Sarstedt) containing PBS (Sigma) with 2% fetal bovine serum (FCS) (Biowest). Red blood cells were removed using BD Pharm Lyse (BD Biosciences). Cells were resuspended in PBS/FCS and naïve CD4^+^ T cells were isolated using the naïve CD4^+^ T cell isolation kit (mouse) from Miltenyi Biotec according to their standard protocol. For some experiments the cells were further FACS-sorted using a SONY SH800 cell sorter.

### 2.5. Th1 differentiation culture

Naïve CD4^+^ T cells were isolated from WT, HDAC1^On^ and HDAC1^Off^ mice (8-12 weeks of age) as described above and cultured in a 48-well plate (300000 cells/well) (Sarstedt) in T cell medium supplemented with 20 U/ml rhIL-2 (Peprotech), 5 ng/ml IL-12 (R&D Systems) and 3 μg/ml anti-IL-4 in the presence of plate bound anti-CD3ε (1 μg/ml) (BD Biosciences) and anti-CD28 (3 μg/ml) (BD Biosciences). After 72h, cells were harvested and stained with Fixable Viability Dye eFluor 506 (Thermo Fisher Scientific) and appropriate antibodies and subjected to flow cytometric analysis.

### 2.6. Flow Cytometric analysis

Cells were incubated with Fc-block (1:250; BD Biosciences) followed by surface staining with the appropriate antibodies. Dead cells were excluded using Fixable Viability Dye eFluor 506 or 780 (Thermo Fisher Scientific) according to the manufacturer’s protocol. For the detection of intracellular transcription factor and cytokine expression, cells were fixed and permeabilized using the Foxp3 Staining Buffer Set (Thermo Fisher Scientific) or the BD Cytofix/Cytoperm Fixation/Permeabilization solution kit (BD Biosciences) according to the manufacturer’s protocols and stained with the appropriate antibodies. For some experiments, cells were additionally labelled with CellTrace Violet Proliferation dye (Thermo Fisher Scientific). After staining, cells were measured using a BD FACS Fortessa (BD Biosciences) or a CytoFLEX S (Beckman Coulter) and analyzed using FlowJo v10.2 software (TreeStar).

### 2.7. Flow cytometry antibodies

The following anti-mouse antibodies were used for flow cytometry: CD19 (clone: 6D5), CD4 (clone: RM4-5), CD44 (clone: IM7), CD45.1 (clone: A20), CD45.2 (clone: 104), CD62L (clone: MEL-14), IFNγ (clone: XMG1.2), IL-17A (clone: TC11-18H10.1), CD11a (clone: M17/4) CD18 (clone: M18/2), Vα3.2 (clone: RR3-16), Vβ11 (clone: KT11), all from Biolegend; CD8 (clone: 53-6.7) CD162 (clone: 2PH1), CD43 (clone: S7), all from BD Biosciences. Ki67 (clone: SolA15), TCRβ (clone: H57-597), CD69 (clone: H1.2F3), all from Thermo Fisher Scientific.

### 2.8. Protein isolation

Cell pellets were resuspended in <u>Karin</u> lysis <u>buffer</u> (20 mM Tris-HCl pH 8.0, 138 mM NaCl, 10 mM EDTA, 100 mM NaF, 1% Nonidet P-40, 10% glycerol, 2 mM sodium vanadate) supplemented with cOmplete protease inhibitor cocktail (Roche), 100 μM PMSF, 10 mM sodium fluoride, 10 mM β-glycerophosphate, 10 μM sodium molybdate and 100 μM orthovanadate. Following three freeze-and-thaw cycles and full speed centrifugation, the supernatant containing extracted soluble protein was collected and protein concentration was determined using Bio-Rad Protein Assay Dye Reagent Concentrate (5000006, Bio-Rad) according to the standard protocol provided by the manufacturer.

### 2.9. Immunoprecipitation

For FLAG immunoprecipitation (FLAG-IP), 30 µl of Anti-FLAG M2 Magnetic Beads (M8823, Sigma-Aldrich) were used and after 3 washes in Karin buffer including proteinase inhibitors, beads were blocked for 30 min in 1 mg/ml BSA (Karin buffer including proteinase inhibitors). The beads were incubated overnight with 200 µg - 1 mg of proteins in Karin buffer including proteinase inhibitors at 4°C. The next day, the beads were washed 3× with Karin buffer including proteinase inhibitors. After the last washing step, 2/3 of the beads were used for Western blotting and 1/3 of the beads were used for the histone deacetylase assay. For all immunoprecipitations, background controls with the corresponding amounts of extract were included.

### 2.10. Immunoblot analysis

In case of input samples, 20 µg of whole cellular protein were denatured in sodium dodecyl sulfate (SDS) loading dye. For IP samples, the buffer was completely removed and proteins were eluted from the beads with SDS loading dye (without Dithiothreitol (DTT)) for 5 min at 55°C. The eluate was mixed with SDS loading dye (with DTT) and denatured. Proteins were separated by SDS-polyacrylamide gel electrophoresis and transferred onto a nitrocellulose membrane (Amersham Protran, GE10600001, Sigma Aldrich) by the wet transfer method. The membrane was washed in blocking solution (1× PBS, 1% polyvinylpyrrolidone, 1% non-fat dried milk, 0.1% tween-20, 0.01% sodium azide, pH 7.4) and incubated with the following primary antibodies: FLAG M2 (F1804, Sigma), HDAC1 (Sat208, Seiser Lab or 10E2 Seiser/Ogris Lab), HDAC2 (ab7029, Abcam or 3F3, Seiser/Ogris Lab), SIN3A (sc994, Santa Cruz or ab3479, Abcam), CoREST (RCOR1, 07-455, Millipore), MTA2 (276, Sigma or M7569, Sigma) and β-actin (ab8226, Abcam or ab8227, Abcam). For detection, the ECL Western blotting detection reagents (RPN2106, GE Healthcare) were used together with a FUSION FX chemiluminescence imaging system.

### 2.11. Deacetylase activity assay using histones

The deacetylase assay was conducted in 20 µl total volume containing beads with immunoprecipitated proteins and 4 µl of [^3^H]-acetate-labeled chicken erythrocyte histones (1.5-2 mg/ml) as described previously [34]. Following incubation for 5 hours on a thermomixer (300 rpm, 30°C), 35 μl of Histone Stop Solution (1 M HCl, 0.4 M sodium acetate) and 800 μl of ethyl acetate were added. After vortexing for 15 sec, the samples were centrifuged with a swing-out bucket centrifuge (10000 rpm, 4 min, RT). 600 µl of the organic phase were mixed with 3 ml of Scintillation Solution (5 g/l PPO, 0.5 g/l POPOP, in toluene). Released [^3^H]-acetate (counts per minute) was determined with a Liquid Scintillation Analyzer (Packard).

### 2.12. Cycloheximide chase assay

To determine the stability of HDAC1^On^ and HDAC1^Off^ proteins, naïve CD4^+^ T cells were activated overnight in a 48-well plate (1×10^6^ cells/well) (Sarstedt) in T cell medium in the presence of plate bound anti-CD3ε (1 μg/ml) (BD Biosciences) and anti-CD28 (3 μg/ml) (BD Biosciences). After activation, cells were treated with 300 µg/ml Cycloheximide (Sigma). After the indicated time points, cells were harvested, stained with Fixable Viability Dye eFluor 506 (Thermo Fisher Scientific), anti-FLAG and anti-CD4 and subjected to flow cytometric analysis.

### 2.13. Experimental autoimmune encephalomyelitis (EAE)

To induce EAE, female 2D2-HDAC1^On^ and 2D2-HDAC1^Off^ mice were subcutaneously (s.c.) injected into the left and right flank with 100 μl of an emulsion of MOG_35-55_ peptide (500 μg/ml) (M4939, Sigma Aldrich) in distilled water and complete Freund’s adjuvant (made from incomplete Freund’s adjuvant (Sigma) that has been supplemented with 5 mg/ml heat-killed Mycobacterium tuberculosis (strain H37Ra; Difco)). On the day of immunization as well as 2 days later mice were injected intraperitoneally (i.p.) with 400 ng pertussis toxin (Sigma Aldrich). Disease progression in mice was monitored daily and assessed according to the following disease scores: 0 = unaffected; 0.5 = incomplete limp tail; 1 = complete limp tail; 1.5 = limp tail and hind limb weakness (mild wobbly walk); 2 = limp tail and weakness of hind legs (wobbly walk) 2.5 = limp tail and dragging of hind legs; 3 = no movement or complete dragging of one leg; 3.5 = complete bilateral paralysis of hind legs; 4 = limp tail and complete paralysis of hind legs, or mouse is unable to turn upright when placed on its side, or mouse is dragging the complete hindquarter; 4.5 = limp tail, complete hind leg and partial front leg paralysis but mouse is alert and feeding; 5 = complete hind and partial front leg paralysis, no movement.

### 2.14. Low-input RNA sequencing of in vivo activated CD4^+^2D2^+^ T cells

6 × 10^5^ naïve 2D2-HDAC^On^ or 2D2-HDAC1^Off^ CD4^+^ T cells (CD45.2^+^) were transferred (i.v.) into CD45.1^+^ recipient mice. 18 h after cell transfer recipient mice were immunized into the right hind paw with 50 μl of a 1:1 emulsion of 1 mg/ml MOG_35-55_ peptide (M4939, Sigma Aldrich) in distilled water and complete Freund’s adjuvant (incomplete Freund’s adjuvant (Sigma) supplemented with 10 mg/ml heat-killed Mycobacterium tuberculosis (strain H37Ra; Difco)). Draining (popliteal) lymph nodes were isolated 4 days later and single cell suspensions were prepared. Two mice were pooled for one biological replicate and three biological replicates were generated for each genotype. Between 10,000 and 15,000 CD45.2^+^ 2D2-HDAC1^On^ and 2D2-HDAC1^Off^ CD4^+^ T cells were sorted. Total RNA was prepared from cell lysates using the RNeasy Mini Kit (Qiagen) and RNase-Free DNase Set (Qiagen) according to manufacturer’s protocol. RNA and library concentrations were determined using Qbit 2.0 Fluorometric Quantitation (Life Technologies). RNA and library integrities were determined using Experion Automated Electrophoresis System (Bio-Rad). Library preparation and RNA Sequencing were performed by the Biomedical Sequencing facility at CeMM (Research Center for Molecular Medicine Vienna, Austria) using low-input Smart-seq2 [35]. The libraries were sequenced using the Illumina HiSeq 3000 platform and the 50-bp single-read configuration.

### 2.15. Bioinformatic analysis of RNA-seq data

Raw sequencing data were processed with Illumina2 bam-tools 1.17 to generate sample-specific, unaligned BAM files. Sequence reads were mapped onto the mouse genome assembly build mm10 (a flavor of GRCm38) using TopHat 2.0.13 [36]. Gene expression values (reads per kilobase exon per million mapped reads) were calculated with Cufflinks 2.2.1 [37]. The statistical analysis was performed in R. Raw count matrices were filtered for a minimum of 10 counts in all samples to account for outlier genes and mitigate the overall high variance in the data. Differential expression testing was performed using the Bioconductor package DESeq2 (v1.34.0) [38]. Fold change shrinkage was applied in order to moderate estimated fold changes of low count genes. Gene Set Enrichment Analysis (GSEA) was performed using the Bioconductor package clusterProfiler (v4.2.2) [39] and visualized using functions from the Bioconductor package enrichplot (v1.14.2) [40]. Other data was visualized using the ggplot2 package [41].

### 2.16. Cell extract and mass spectrometry sample preparation

Cells were lysed in Karin lysis buffer supplemented with cOmplete protease inhibitor cocktail (Roche), 100 μM PMSF, 10 mM sodium fluoride, 10 mM β-glycerophosphate, 10 μM sodium molybdate and 100 μM orthovanadate, 5 mM sodium butyrate and 10ng/ml Nicotinamide (NAM). 250 U of benzonase (Merck) were added to each sample. Lysis was achieved using a Bioruptor sonication device (Diagenode) applying 5 alternating cycles of 30 sec sonication and 30 sec cooling. Power level was set to H. Following lysis, cells were centrifuged at 15,000 × *g* for 10 min at 4°C to remove insoluble material. For purification of HDAC1-Flag 1 mg of protein was mixed with 120 µl of crosslinked M2-Flag beads (Sigma Aldrich, M8823-5ml) and incubated rotating overnight at 4°C. The beads were washed three times with TBS, re-buffered to and washed once with ammonium bicarbonate (ABC) (Sigma Aldrich, 09830-500G) buffer, pelleted by gentle centrifugation, and resuspended in 50 µl 2 M urea (VWR, 0568-500G), 50 mM ABC. Pre-digestion of bound proteins was performed by addition of 200 ng / µl LysC (FUJIFILM Wako Pure Chemical Corporation, 125-02543) and trypsin (Trypsin Gold, Mass Spec Grade, Promega, V5280) mixed 1:1, followed by an incubation at RT for 90 min in the dark. Pre-digested proteins and peptides were eluted from the beads, and beads were subsequently rinsed with 50 µl ABC and pelleted by gentle centrifugation. The eluates were united. Pre-digested proteins and peptides were reduced with 10 mM dithiothreitol (DTT) (Roche, 10197777001) for 30 min at RT, and subsequently carbamidomethylated with 20 mM iodoacetamide (IAA) (Sigma Aldrich, I6125-5G) for 30 min at RT in the dark. The alkylation reaction was quenched by adding 5 mM DTT for 10 min. Samples were digested overnight at 37°C using 150 ng / µl of the 1:1 LysC and trypsin mix. Following digestion, samples were acidified using 10% trifluoroacetic acid (TFA) (Thermo Scientific, 28903) to a final concentration of 0.5% and desalted on C18 (Empore, 2215-C18) StageTips as described in [42].

### 2.17. Mass spectrometry measurements

Mass spectrometry measurements were performed on an Orbitrap Exploris 480 mass spectrometer (Thermo Fisher). The instruments were operated in data-dependent acquisition (DDA) mode with dynamic exclusion enabled. For peptide separation on the HPLC, the concentration of organic solvent (acetonitrile, VWR, 83639.320) was increased from 1.6% to 28% in 0.1% formic acid at a flow rate of 230 nl / min, using a 2 hours gradient time. Peptides were ionized with a spray voltage of 2.3 kV. MS1 survey scans were recorded with the following parameters: resolution 60,000, scan range 350-1,500 m/z, automatic gain control (AGC) target = Custom, and maximum injection time (IT) mode = Auto, FAIMS voltages on, FAIMS CV -45V and -60V, dynamic exclusion was set to 45 sec. MS2 analysis (CID) was performed with the following parameters: resolution 15,000, normalized AGC Target 100%, AGC target = Custom, maximum IT mode = Custom, isolation window 1.2 m/z, and HCD collision energy 34%.

### 2.18. Mass spectrometry data analysis

Mass spectrometry raw files were searched using MaxQuant (version 2.0.3.0) against the *Mus Musculus* (mouse) Uniprot database (release 2022.01, FASTA file: 2022.01_UP000000589_10090_Mus_musculus_1protein_per_gene.fast) and a database of common laboratory contaminants (provided with MaxQuant). Enzyme specificity was set to “Trypsin/P”. Carbamidomethylation of cysteine was searched as a fixed modification. Protein N-term acetylation and Oxidation (M) were set as variable modifications. A maximum of 5 variable modifications per peptide was allowed. Minimal peptide length was set to 7, the maximum number of missed cleavages was set to 2, and all data were filtered at 1% FDR (PSM and protein). Further data processing was performed using the Cassiopeia script [43] Reverse hits, contaminants and proteins with less than 2 razor & unique peptides were removed for the quantitative analysis. Missing label free quantitation (LFQ) values were imputed after logarithmic transformation by drawing random values from a normal distribution (estimated from sample LFQ intensity distribution and shifted by -1.8 standard deviations with a width of 0.3 standard deviations). For pairwise comparison the log2 fold change and mean LFQ were calculated and plotted against each other for all hits with more than 3 MSMS counts (MAplot).

### 2.19. Statistical analysis

No statistical methods were used to predetermine the sample size. All statistical analyses were performed using Prism 9 Software (GraphPad Inc). As indicated in each figure legend, p-values were calculated using either unpaired two-tailed t-test, one-way or two-way ANOVA. No data were excluded and no specific randomization of animals or blinding of investigators was applied. The data shown indicate the mean.

### 2.20. Protein network analysis

Protein-protein interaction networks were obtained using the STRING database v11.5 [44]. Network parameters were exported from STRING DB as short tabular text output and modified: column “node1” was defined as “source”, column node 2 was defined as “target” and the combined STRING score was used to define the weight of the network edge. The data was imported into Gephi v0.9.7 [45] for the network visualization, using the Fruchteman Reingold function with default settings and ForceAtlas 2 function with close to similar scaling and gravity settings, and the “prevent overlaps” option being activated. Node size was set proportional to the degree value and edge thickness to the combined interaction score. Nodes were colored according classification presented in [46].

### 2.21. Data availability statement

Mass spectrometry raw data have been deposited to the ProteomeXchange Consortium (http://proteomecentral.proteomexchange.org) via the PRIDE partner repository [47] with the dataset identifier PXD040347. RNA-seq data reported in this study have been deposited in the National Center for Biotechnology Information’s Gene Expression Omnibus database under number GSE228259.

## 3. Results

### 3.1. Generation of a conditional mouse model for HDAC1 inactivation

To investigate the effect of HDAC1 enzymatic inactivation in contrast to HDAC1 deletion, we created a mouse model conditionally replacing the expression of endogenous HDAC1 by a transgenic version of either catalytically inactive (CI) HDAC1 or, as a control, wildtype (WT) HDAC1 enzyme. Open reading frames for the respective HDAC1 isoforms were cloned into the PBTV2-11 exchange vector and integrated into the targeting platform at the murine *Rosa26* locus *via* ΦC31 integrase mediated recombination (Fig. 1A). Transgenic HDAC1 mRNA and proteins are expressed upon Cre-mediated deletion of a STOP cassette (Fig. 1A). An IRES-EGFP reporter allows to monitor the efficient removal of the STOP cassette. The catalytically inactive version of HDAC1 carries a single histidine residue to alanine exchange in the catalytic center (H141A) and was previously shown to have strongly reduced deacetylase activity [31, 48–50]. The encoded transgenic catalytically inactive HDAC1 H141A enzyme (hereafter referred to as HDAC1^Off^) and the corresponding transgenic HDAC1 WT enzyme (hereafter referred to as HDAC1^On^) carry a C-terminal FLAG tag, allowing the detection, isolation and quantification of these proteins *via* immunoblot, immunoprecipitation (IP) and flow cytometry analysis. By intercrossing the transgenic *Rosa26* knock-in mouse model with the previously described T cell-specific conditional HDAC1 knockout strain (*Hdac1*^f/f^ *Cd4*-Cre) [30] we ablated endogenous HDAC1 and simultaneously induced the expression of transgenic HDAC1^Off^ or HDAC1^On^ protein in T cells. A schematic description of the mouse genotypes used in this study are shown in Fig. 1B. Analyzed mice were either WT (*Hdac1^fl/fl^ Cd4-Cre^–^*), HDAC1^cKO^ (*Hdac1^fl/fl^ Cd4-Cre^+^*), HDAC1^On^ (*Hdac1^fl/fl^ Rosa26-STOP^fl^-HDAC1^WT^ Cd4-Cre^+^*) or HDAC1^Off^ (*HDAC1^fl/fl^ Cd4*-Cre^+^ *Rosa26-STOP^fl^-HDAC1^Off^ Cd4-Cre^+^*).

In order to confirm that the generated HDAC1^Off^ mice are indeed suitable to study our translational research question, we first determined the expression levels of the two HDAC1 versions driven from the *Rosa26* locus. Naïve CD4^+^ T cells were isolated from HDAC1^Off^ and HDAC1^On^ mice and activated with anti-CD3/anti-CD28 in Th1 polarizing conditions. Subsequently, protein extracts were isolated and an immunoblot analysis revealed that HDAC1^Off^ and HDAC1^On^ proteins were expressed at similar levels (Fig. 1C). Moreover, over 90% of *in vitro* differentiated T cells expressed FLAG-tagged HDAC1^Off^ or HDAC1^On^ protein as assessed by flow cytometry (Figs. 1D and 1E). Consistent with the detection of FLAG-tagged HDAC1 forms, we also observed expression of EGFP (via the IRES-EGFP cassette) in more than 90% of activated CD4^+^ T cells (Fig. 1F). These data show that we successfully generated two novel mouse strains expressing either HDAC1^Off^ or HDAC1^On^ proteins from the *Rosa26* locus in T cells while at the same time lacking endogenous HDAC1.

### 3.2. HDAC1^Off^ is stable and exhibits decreased deacetylase activity compared to HDAC1^On^

Catalytically inactive mutant isoforms of several enzymes have altered protein stability [51–54]. Therefore, we examined whether the HDAC1^Off^ protein exhibits a similar or an altered stability compared to the HDAC1^On^ protein by performing cycloheximide time course experiments. Cycloheximide is an inhibitor of protein biosynthesis and is commonly used to determine the half-life of a given protein [55]. Naïve HDAC1^Off^ and HDAC1^On^ CD4^+^ T cells were activated overnight using anti-CD3/anti-CD28 and then HDAC1^On^ and HDAC1^Off^ protein stability was assessed in the course of 24 hours cycloheximide treatment by flow cytometry (Figs. 1G and 1H). There was no difference in the abundance (measured as mean fluorescence intensity) between HDAC1^On^ and HDAC1^Off^ proteins after 2, 4 and 6 hours of cycloheximide treatment. There was a slight tendency of HDAC1^Off^ protein being more abundant after 8, 16 and 24 hours compared to the HDAC1^On^ version, however this difference did not reach statistical significance. These data indicate that loss of catalytic activity does not have a major impact on the stability of HDAC1 in activated CD4^+^ T cells.

Next, we analyzed and compared the catalytic activity of HDAC1^Off^ and HDAC1^On^ proteins in T cells. Again, naïve CD4^+^ T cells were isolated from HDAC1^Off^ and HDAC1^On^ mice and activated under Th1 polarizing conditions. Protein extracts from activated HDAC1^Off^ and HDAC1^On^ CD4^+^ T cells were isolated, subjected to a FLAG-immunoprecipitation (FLAG-IP) followed by a total deacetylase activity assay with ^3^H-acetylated histones as substrate (Fig. 1I) [34, 56]. As expected, we identified a strong and significant reduction in the deacetylation activity of precipitated HDAC1^Off^ protein compared to HDAC1^On^ protein (Fig. 1I, right panel). As negative controls, we used protein extracts from WT and HDAC1^cKO^ Th1 cells lacking FLAG-tagged HDAC1 protein. Of note, we observed a slight decrease in the deacetylase activity of HDAC1^Off^ whole cell input extracts (Fig. 1I, left panel). This indicates that HDAC1 is a major HDAC isoform contributing to the total deacetylase activity in Th1 cells and that other HDAC members are not able to fully compensate the loss of HDAC1 catalytic activity. These data show that a histidine 141 to alanine mutation strongly reduces the catalytic activity of HDAC1 without significant effects on its protein stability in CD4^+^ T cells.

Taken together, the characterization of CD4^+^ T cells isolated from HDAC1^Off^ and HDAC1^On^ mice indicates that we have generated novel mouse models that allow us to study the impact of the inactivation of HDAC1 enzymatic activity in *in vitro* and *in vivo* settings. Of note, non-immunized HDAC1^Off^ and HDAC1^On^ mice had similar frequencies and numbers of CD4^+^ and CD8^+^ T cells as well as B cells in the spleen (Supplementary Fig. 1A.). Furthermore, HDAC1^Off^ and HDAC1^On^ splenic CD4^+^ T cells also showed similar IFNγ and IL-17A expression upon *ex vivo* PMA/ionomycin stimulation (Supplementary Fig. 1B). This indicates that the expression of HDAC1^Off^ protein in T cells does not lead to major alterations of T cell subset composition at steady state in HDAC1^Off^ mice in comparison to the corresponding HDAC1^On^ mice.

### 3.3. HDAC1^Off^ and HDAC1^On^ proteins are incorporated into co-repressor complexes and have similar interactomes

It was previously shown that HDAC1 function depends on its proper incorporation into co-repressor complexes, including SIN3A, NuRD [57] and COREST (also known as RCOR1) [10, 57–62]. We therefore investigated whether the catalytic activity is required for HDAC1 to be incorporated into co-repressor complexes. Protein extracts from *in vitro* differentiated HDAC1^Off^ and HDAC1^On^ Th1 cells were generated and subjected to FLAG-IP followed by Western blot analysis. This analysis showed that comparable amounts of components of the co-repressor complexes SIN3A, NuRD (using MTA2 as component) and COREST co-precipitated with HDAC1^Off^ and HDAC1^On^ (Fig. 2A). We additionally examined whether HDAC2 co-precipitates with the HDAC1^Off^ protein to a similar extent as with the HDAC1^On^ protein, knowing that HDAC1 and HDAC2 interact and form the catalytic core of several co-repressor complexes [9, 10]. Indeed, FLAG-IP experiments revealed that HDAC2 co-precipitated with both HDAC1^Off^ and HDAC1^On^ proteins at similar levels (Fig. 2A). This data indicates that the catalytic activity of HDAC1 is not required for its association with HDAC2 and components of co-repressor complexes.

**Figure 2.**
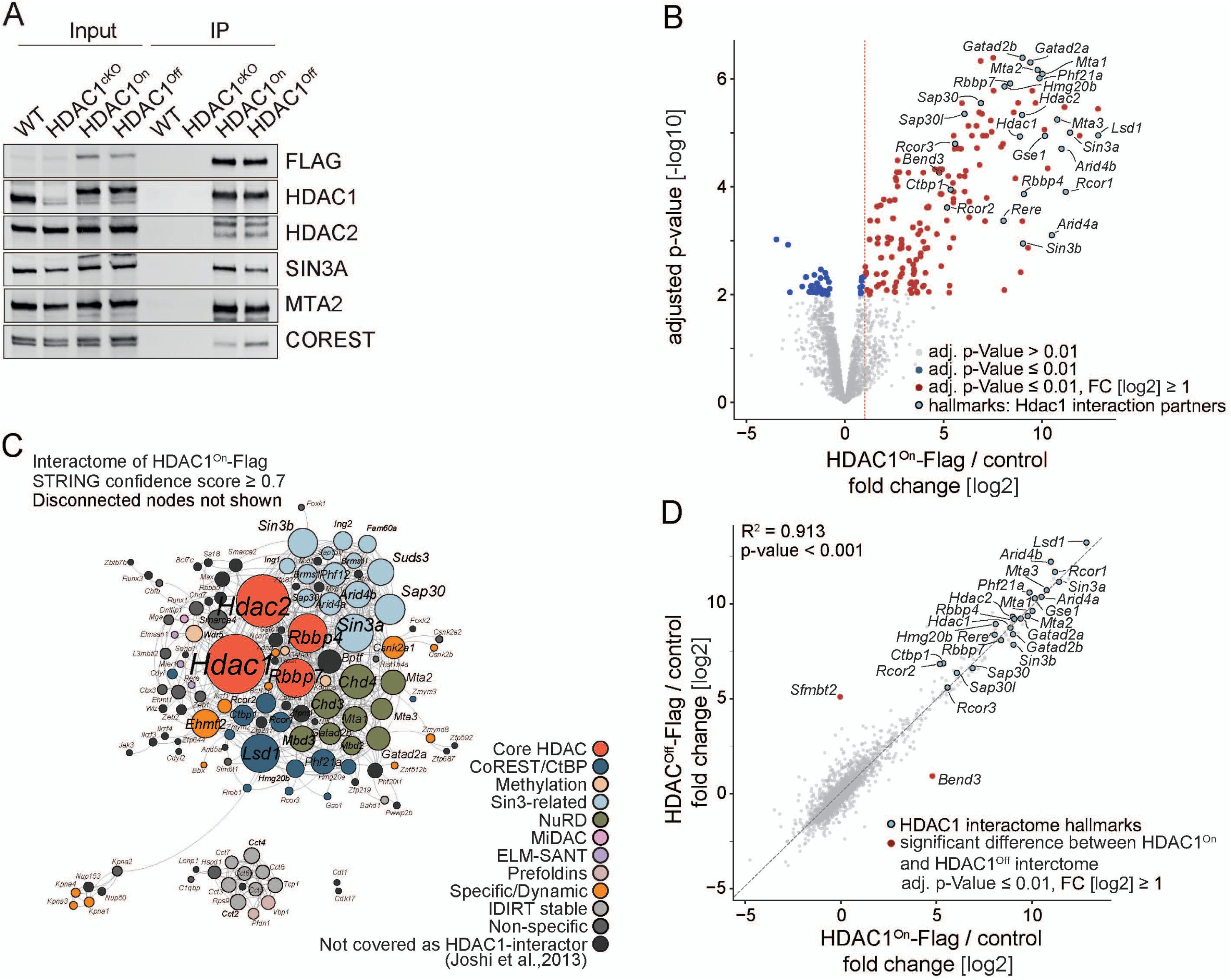
HDAC1^On^ and HDAC1^Off^ proteins have similar interactomes in CD4^+^ T cells. **(A)** Western blot analysis of total cell extracts (Input, left panel) and of FLAG immunoprecipitates (IP, right panel) isolated from WT, HDAC1^cKO^, HDAC1^On^ and HDAC1^Off^ Th1 cells using antibodies specific for the FLAG epitope, HDAC1, HDAC2, SIN3A, MTA2 or COREST. Data shown are representative of 3 independent experiments. **(B)** Volcano plot highlighting the interactome of HDAC1^On^ protein as determined by mass spectrometry. X-axis shows fold changes in abundance of HDAC1^On^-associated proteins, expressed as mean LFQ intensities [log2] of HDAC1^On^ FLAG-IP over the Cre-negative control. Y-axis: Adjusted p values [-log10]. Proteins showing a fold change ≥ 2-fold ([log2] >1) and an adjusted p value ≤ 0.01 were considered as interactors of HDAC1^On^. **(C)** STRING DB-based protein network analysis of the HDAC1^On^ interactome identified in (B). Color coding is according to reference [46] **(D)** Scatter plot comparing global fold changes (FLAG-IP over control [log2]) of interacting proteins of HDAC1^Off^ to HDAC1^On^. Examples of well-known members of co-repressor complexes are indicated in light blue. Proteins which are exclusively part of the HDAC1^Off^ or HDAC1^On^ interactome are indicated in red. (B-D) The gene names of interacting proteins are shown.

To further examine whether the inactivation of the catalytically activity affects the complex composition of HDAC1-associated proteins, we analyzed global interactomes of C-terminally FLAG-tagged HDAC1^On^ and HDAC1^Off^ in Th1 cells using affinity purification mass spectrometry (AP-MS). We identified 167 proteins (FC ≥ 2-fold, adjusted p-value ≤ 0.01 compared to the negative Cre^–^ control) as potential interaction partners of HDAC1^On^ (Fig. 2B, Supplementary Table 1). This set of putative interactors covered the well-known subunits of co-repressor complexes as also revealed by co-immunoprecipitation (Fig. 2A), such as CoREST, Sin3-related and NuRD (Fig. 2B and 2C). Moreover, the set of interactors was identical for both HDAC1 isoforms and we identified only two proteins being differentially bound between HDAC1^Off^ and HDAC1^On^, namely the transcriptional repressor proteins Bend3 and Sfmbt2 (Fig. 2D). Our results therefore indicate that loss of the catalytic activity of HDAC1 does not affect the HDAC1 interactome.

### 3.4. HDAC1^Off^ mice are protected from experimental autoimmune encephalomyelitis

We have previously shown that a conditional deletion of HDAC1 in T cells protects mice against the development of EAE [16, 26]. However, it remains elusive whether the specific inhibition of HDAC1 catalytic activity in CD4^+^ T cells is sufficient to protect mice against EAE. This is a key issue to further the translational potential and therapeutic concept of targeting HDAC1. To address this question, we induced EAE by immunizing HDAC1^Off^ and HDAC1^On^ mice with myelin oligodendrocyte glycoprotein (MOG) peptide_35-55_/CFA on day 0 followed by pertussis toxin injection on day 0 and day 2, as previously described [26]. As expected, all HDAC1^On^ mice (n=10, analyzed in two independent experiments with 5 mice each) developed clinical signs of EAE and thus displayed a 100% disease incidence. HDAC1^On^ mice showed a strong disease induction around day 11 with a peak in the clinical EAE score around day 18 (disease score approx. 2.5) (Fig. 3A). In contrast, 9 out of 10 HDAC1^Off^ mice (analyzed in two independent experiments with 5 mice each) developed no clinical EAE symptoms (clinical score 0), while 1 HDAC1^Off^ mouse developed EAE symptoms with a clinical score of 2. This resulted in an overall clinical EAE score of 0.2 across all HDAC1^Off^ mice (Fig. 3A). A detailed analysis of the CNS of HDAC1^On^ mice at the peak of disease (day 18) revealed a strong infiltration of leukocytes (Fig. 3B), while the total number of leukocytes isolated from the CNS of HDAC1^Off^ mice was severely reduced (Fig. 3B). Although frequencies of B cells as well as of CD4^+^ and CD8^+^ T cells among the CNS infiltrating cells were not altered in HDAC1^Off^ mice compared to HDAC1^On^ mice (Fig. 3C and 3D), there was a tendency that the number of CNS-infiltrating HDAC1^Off^ CD4^+^ T cells was reduced in comparison to infiltrating HDAC1^On^ CD4^+^ T cells (Fig. 3D). Moreover, CNS CD4^+^ T cells in HDAC1^Off^ mice showed a severe reduction in the expression of IFNγ, IL-17A and TNFα (Fig. 3E and 3F). CD4^+^ T cell numbers in the spleen and draining lymph nodes were slightly enhanced in HDAC1^Off^ mice compared to HDAC1^On^ mice, most likely due to an impaired infiltration of HDAC1^Off^ effector cells into the CNS, although the difference did not reach statistical significance (Supplementary Fig. 2A). We also examined the proliferation state of effector CD4^+^ T cells in the CNS, draining lymph node and spleen by measuring expression of the proliferation marker Ki67 (Supplementary Fig. 2B and 2C). We observed that the few CD4^+^ T cells present in the CNS of HDAC1^Off^ mice showed lower expression of Ki67, suggesting reduced proliferation of effector CD4^+^ T cells. In contrast, Ki67 expression in dLN and splenic CD4^+^ T cells was similar between HDAC1^Off^ and HDAC1^On^ mice. Together, these data indicate that the inhibition of HDAC1 enzymatic activity results in a reduced infiltration of leukocytes and recruitment of CD4^+^ T cells into the CNS and impaired effector function of CD4^+^ T cells and as a consequence absence of clinical EAE signs.

**Figure 3.**
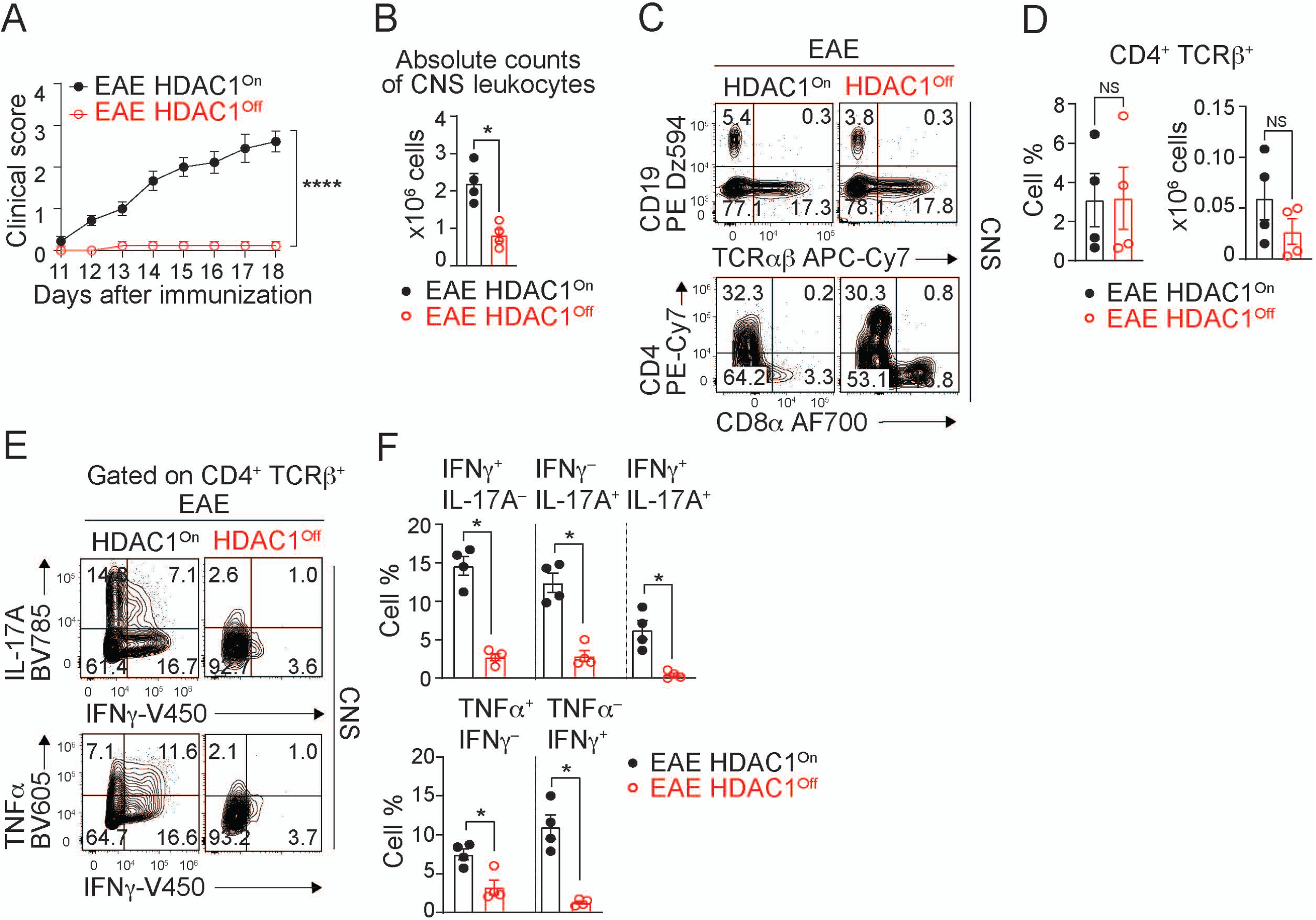
HDAC1^Off^ mice do not develop experimental autoimmune encephalomyelitis. **(A)** Diagram showing the EAE clinical score of HDAC1^On^ and HDAC1^Off^ mice at different days following disease induction with MOG_35-55_ peptide in CFA. **(B)** Total leukocyte cell numbers in the CNS around the peak of disease (day 18, as determined with HDAC1^On^ mice). **(C)** Flow cytometry analysis showing CD19, TCRβ, CD4 and CD8 expression among CNS infiltrating leukocytes. (**D**) Diagrams show the percentage and total cell numbers of CD4^+^ T cells. **(E)** Cytokine expression profiles (IL-17A, IFNγ and TNFα) of CNS CD4^+^ T cells isolated on day 18. Cells were restimulated with PMA/ionomycin. (**F**) Diagrams show the percentage of CD4^+^ T cells that express the indicated cytokines. Data in (A) show mean ± SEM, two-way analysis of variance (ANOVA). (B,D,F) Bar diagrams show mean ± SEM; Student t-test (unpaired, two-sided with Welch’s correction). (A,D) *p≤0.05, ****p≤0.0001. Data show the summary (A,B,D,F) or a representative (C,E) of 10 HDAC1^On^ and 10 HDAC1^Off^ mice analyzed in 2 experiments (A) or of 4 HDAC1^On^ and 4 HDAC1^Off^ mice/CD4^+^ T cells analyzed in 1 experimental round (B-F).

### 3.5. Impact of HDAC1 catalytic activity inactivation on gene expression

Having shown that the inhibition of the catalytic activity of HDAC1 in T cells is sufficient to protect mice from EAE, we next wanted to understand the impact of HDAC1^Off^ and HDAC1^On^ expression on the transcriptomes of *in vivo* activated antigen-specific CD4^+^ T cells. In order to do so, we followed a protocol that we previously applied to investigate transcriptomes of *in vivo* activated WT and HDAC1^cKO^ CD4^+^ T cells [27]. We crossed HDAC1^On^ and HDAC1^Off^ mice with 2D2 TCR transgenic mice that express a TCR (formed by Vα3.2 and Vβ11) specific for MOG_35-55_ peptide [29]. We transferred either 2D2-HDAC1^On^ or 2D2-HDAC1^Off^ CD4^+^ T cells (CD45.2^+^) into CD45.1^+^ mice and immunized recipient mice with MOG_35-55_/CFA. On day 4 post-immunization, 2D2 CD4^+^ T cells were isolated from the draining lymph nodes and subjected to low-input Smart-seq2 RNA-sequencing. We detected 181 genes being upregulated and 178 genes being downregulated in HDAC1^Off^ CD4^+^ T cells compared to HDAC1^On^ CD4^+^ T cells (Fig. 4A and Supplementary Table 2). To identify pathways and hallmark gene sets that are altered in HDAC1^Off^ T cells, we performed gene set enrichment analysis (GSEA) using the Gene Ontology (GO) database. Several gene sets (GO terms: biological processes; GO:BP) were underrepresented in HDAC1^Off^ compared to HDAC1^On^ cells, including pathways that play a role in cell motility and migration including “positive regulation of cell migration”, “positive regulation of cell motility” and “cell adhesion” (Fig. 4B). Moreover, pathways related to T cell effector function such as “response to cytokine”, “cytokine production” and “T cell activation” were reduced in HDAC1^Off^ CD4^+^ T cells. A list of all identified hallmark GO:BP gene sets from this analysis can be found in Supplementary Table 3. Thus, the RNA-seq data showed that *in vivo*-activated CD4^+^ T cells only expressing catalytically inactive HDAC1 are unable to efficiently express genes important for T cell activation and T cell migration in comparison to HDAC1^On^ CD4^+^ T cells.

**Figure 4.**
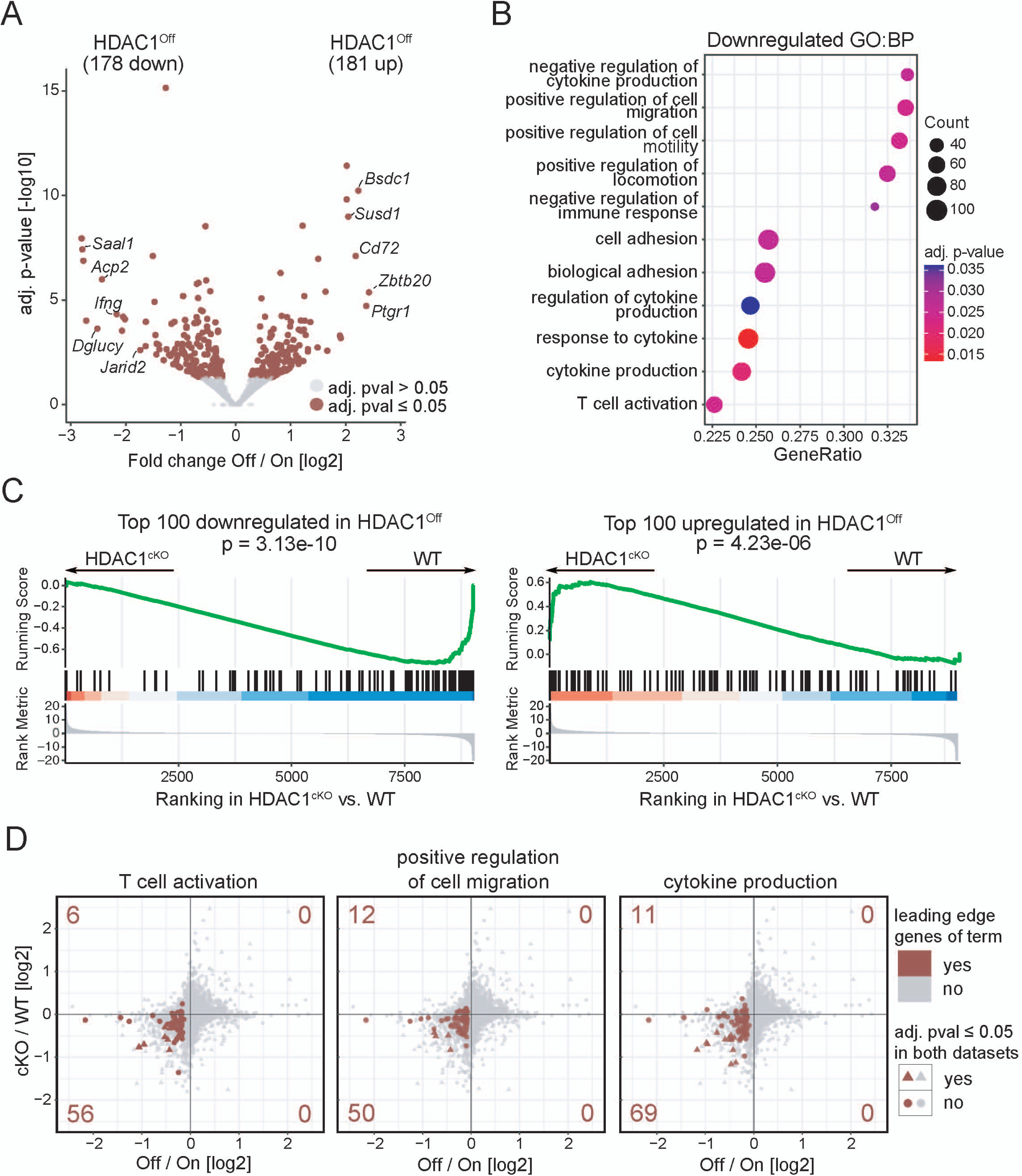
RNA-seq analysis reveals similar transcriptional alterations in *in vivo* activated HDAC1^Off^ and HDAC1^cKO^ CD4^+^ T cells. Naive 2D2-HDAC^On^ or 2D2-HDAC1^Off^ CD4^+^ T cells (CD45.2^+^) were injected into recipient mice (CD45.1^+^) followed by MOG_35-55_/CFA immunization. 4 days later, 2D2^+^CD45.2^+^ CD4^+^ T cells were isolated (from three independent batches) and subjected to low-input Smart-seq2 sequencing. **(A)** Volcano plot depicts the comparison of gene expression profiles between 2D2-HDAC1^On^ and 2D2-HDAC1^Off^ CD4^+^ T cells. On the x-axis the Off/On log2 fold change is plotted, the y-axis indicates FDR-adjusted p-values. 178 genes are downregulated and 181 upregulated in 2D2-HDAC1^Off^ CD4^+^ T cells compared to 2D2-HDAC1^On^ CD4^+^ T cells (FDR ≤ 0.05). The top 5 dysregulated protein-coding genes on each side are labeled with their respective gene symbol. Adjusted p-values were capped at 10^-15^ in the plot. **(B)** Dot plot illustrating results of Gene Set Enrichment Analysis (GSEA) for selected gene sets from the Gene Ontology database term Biological Processes (GO:BP). The x-axis indicates the fraction of genes within each gene set that are part of the leading edge subset. **(C)** GSEA to explore the similarity between HDAC1^Off^ and HDAC1^cKO^ datasets. The top 100 downregulated and the top 100 upregulated genes in (A) were separately tested for enrichment in the entire ranked gene list from the dataset comparing *in vivo* activated 2D2-HDAC1^cKO^ versus 2D2-WT CD4^+^ T cells. **(D)** Scatterplot comparing fold changes of genes in the HDAC1^Off^ dataset (featuring HDAC1^Off^ versus HDAC1^On^) and the HDAC1^cKO^ dataset (featuring HDAC1^cKO^ versus WT). Each data point corresponds to a single gene. The leading edge genes of three enriched gene sets shown in (B) (T cell activation, positive regulation of cell migration and cytokine production) appear in red. The red numbers in each corner denote the corresponding number of leading edge genes in each plot quadrant. Triangular data points represent genes that were found significant in both datasets (FDR ≤ 0.05).

Previous work from our lab has identified that EAE protection in HDAC1^cKO^ mice is caused by a migratory defect of HDAC1-deficient T cells and is accompanied by downregulation of genes associated with leukocyte extravasation [27]. Given that our RNA-seq data indicate a down-regulation of pathways associated with cell migration, and since CD4^+^ T cell trafficking critically mediates immunopathogenicity of EAE [19], we investigated if the expression of certain factors associated with cell migration is altered in HDAC1^Off^ mice, as observed for HDAC1^cKO^ mice. HDAC1^Off^ mice showed reduced expression of the P-selectin ligand CD162 on effector CD4^+^ T cells of spleen while expression levels of LFA-1 integrin chains CD11a and CD18 were not changed (Supplementary Fig. 2D and 2E).

Finally, to study in an unbiased manner whether similar genes are differentially expressed in CD4^+^ T cells upon loss of HDAC1 catalytic activity (HDAC1^Off^ versus HDAC1^On^) or loss of protein (HDAC1^cKO^ versus WT) [27], additional GSEA were performed. For this we compared our HDAC1^Off^ and HDAC1^On^ 2D2 CD4^+^ T cell RNA-seq data and assembled gene sets of the top 100 HDAC1^Off^ up-regulated (“HDAC1^Off^-up”) and the top 100 HDAC1^On^ up-regulated (“HDAC1^On^-up”; i.e. downregulated in HDAC1^Off^) genes (Supplemental Table 2). Subsequently, we assessed enrichment of these two gene sets in our previously generated RNA-seq data from *in vivo* activated WT versus HDAC1^cKO^ 2D2 CD4^+^ T cells [27]. The “HDAC1^Off^-up” gene set was significantly enriched in HDAC1^cKO^ 2D2 CD4^+^ T cells, while the “HDAC1^On^-up” gene set showed a highly significant positive correlation and an enrichment in WT 2D2 CD4^+^ T cells (Figs. 4C and Supplementary Table 4). Upon closer investigation, the enrichment of “HDAC1^Off^-up” and “HDAC1^On^-up” gene sets in HDAC1^cKO^ and WT CD4^+^ T cells, respectively, can be explained by a similar dysregulation of genes implicated in biological processes such as “T cell activation”, “positive regulation of cell migration” and “cytokine production” (Fig 4D). Thus, there is a significant overlap of dysregulated genes and pathways in HDAC1^Off^ and HDAC1^cKO^ CD4^+^ T cells, indicating that the inability of HDAC1-deficient CD4^+^ T cells to induce EAE is due to the loss of HDAC1 catalytic activity.

Taken together, we have generated a novel mouse model to study the impact of T cell specific HDAC1 inactivation and have demonstrated that HDAC1 catalytic activity is crucial for the onset of EAE in mice.

## 4. Discussion

In this study we describe the generation of a novel mouse model allowing the conditional replacement of the expression of endogenous wildtype HDAC1 by a catalytically inactive HDAC1 isoform specifically in T cells. This HDAC1^Off^ model mimics the clinical application of an HDAC1 specific inhibitor and thus made it possible to specifically address the role of the catalytic activity of HDAC1 in EAE disease development. This is a key question, since we have previously demonstrated that conditional deletion of HDAC1 in T cells protects mice from developing EAE [26, 27], raising the exciting possibility that HDAC1 might be a promising target for the treatment of MS. Indeed, our study revealed for the first time that specific targeting of the catalytic activity of HDAC1 is sufficient to achieve a clinical benefit in the protection against a T cell-mediated autoimmune disease.

One important finding of our study, as part of the generation and characterization of the HDAC1^Off^ mouse model, is the observation that catalytically inactive HDAC1 was efficiently and stably expressed in T cells at levels similar to the ones of WT HDAC1. This indicates that the catalytic activity is not essential for the proper regulation of HDAC1 stability. It is known that the function of HDAC1 highly depends on the incorporation into protein complexes [10]. Our characterization of the interactomes of HDAC1^Off^ versus HDAC1^On^ showed that both HDAC1 proteins basically interact with the same set of proteins, including components of co-repressor complexes SIN3, NuRD and CoREST, demonstrating that the catalytic activity of HDAC1 is dispensable for its capacity to interact with other proteins. Thus, the basic biochemical analysis of HDAC1^Off^ mice and CD4^+^ T cells indicates the successful generation of a mouse model to assess non-catalytic and catalytic functions of HDAC1. Of note, we observed a high level of similarity in the protein interaction network of HDAC1 in CD4^+^ T cells with the HDAC1 interactome identified in human HAP1 [63] and CEM-T [46] cell lines (Supplementary Fig. 3). This indicates that our analysis comprehensively covered binding partners of HDAC1 in CD4^+^ T cells. Moreover, this also suggests a high degree of conservation of the HDAC1 interactome among species and different cell types.

A key finding of our study is the observation that the specific targeting of HDAC1 catalytic activity in T cells is sufficient to protect mice from EAE. Our previous results using gene knockout approaches to delete the *Hdac1* gene showed that HDAC1 is an important factor controlling EAE disease development. Thus, our data described in this study clearly demonstrate that the protective effect of HDAC1 deletion is due to the loss of its catalytic activity. Moreover, there was no compensatory upregulation of HDAC2 protein levels in HDAC1^Off^ CD4^+^ T cells as observed in the HDAC1^cKO^ setting, ruling out the possibility that upregulation of HDAC2 contributes to disease protection when HDAC1 is deleted. Recent studies with mice having a T cell-specific deletion of HDAC1 have shown that HDAC1 is key also for driving the development of other autoimmune diseases such as collagen-induced arthritis and adoptive CD4^+^ T cell transfer colitis [16, 27, 64]. Future studies are warranted to address the role of the catalytic activity of HDAC1 also in these disease models and to strengthen the evidence that therapeutic targeting of HDAC1 is a promising strategy to treat autoimmune diseases.

An important issue in the field of HDAC biology addresses non-catalytic functions (i.e. scaffolding functions) of HDACs. The deletion of HDAC1 or the inactivation of the enzymatic activity of HDAC1 results in a similar protective phenotype against EAE induction. This raises then the question whether similar cellular and molecular mechanisms result in the observed phenotype. We have shown that conditional deletion of HDAC1 in T cells compromises their ability to traffic into the CNS [27]. This observation was accompanied by the downregulation of several genes associated with leukocyte extravasation, for example *Itgal*, *Itgb2*, *Selplg* and *Spn*. Based on the transcriptomics analysis of *in vivo* activated HDAC1^On^ and HDAC1^Off^ CD4^+^ T cells, which was performed using the very same experimental setting as the one applied to compare WT and HDAC1^cKO^ CD4^+^ T cells, loss of the catalytic activity seems to have a similar impact on CD4^+^ T cell function as deletion of HDAC1. This was indicated by the downregulation of several pathways associated with cell migration such as “positive regulation of cell migration”, “positive regulation of cell motility” and “cell adhesion”. This is in line with the observed reduced infiltration of leukocytes, including CD4^+^ T cells, into the CNS of HDAC1^Off^ mice. Moreover, the top 100 genes upregulated in either HDAC1^Off^ or in HDAC1^On^ CD4^+^ T cells were enriched either in HDAC1^cKO^ or in WT CD4^+^ T cells, respectively. These findings suggest that transcriptional programs regulating cell migration and hence the ability of CD4^+^ T cells to induce EAE are dependent on HDAC1 catalytic activity. Of note, cytokine production of the few HDAC1^Off^ CD4^+^ T cells that migrated into the CNS was reduced, while cytokine expression was not impaired in infiltrated HDAC1-deficient CD4^+^ T cells [26]. Additional studies are necessary to investigate whether this indicates a non-catalytic function for HDAC1.

Finally, HDAC1 is a key factor driving development and differentiation in many cell lineages and tissues. Our newly generated HDAC1^Off^ and HDAC1^On^ mouse models are a great tool to dissect catalytic and non-catalytic functions in various biological processes, thus facilitating an in-depth understanding of HDAC1 function in many cellular settings. Moreover, HDACs are promising targets for small molecule inhibitors in the treatment of cancer, neurological and immunological disorders. Our mouse models therefore serve as a paradigm for other HDAC family members to mimic the pharmacological treatment with HDAC isoform selective inhibitors.

Taken together, by generating and employing a novel mouse model, our study establishes the translational concept that targeting the catalytic activity of HDAC1 presents a novel therapeutic avenue for treating CD4^+^ T cell-mediated autoimmune disease.

## Authorship contribution statement

C.Z., V.S. and N.S. helped with study design, performed the majority of the experiments, analyzed and interpreted the data, and co-wrote the manuscript. A.H. and A.M. designed and A.H., L.H., V.M. and A.M. generated the conditional catalytically inactive HDAC1 mice. W.E. and C.S. designed and funded the project, interpreted the data, supervised the project, and co-wrote the manuscript. N.S., T.V. and O.A. performed experiments and M.M. performed bioinformatic analyses. W.R. and M.H. performed the mass spectrometry analysis. C.B. supervised the sequencing experiments and associated data analysis. All authors reviewed the manuscript.

## Acknowledgements

The authors thank Michael Schuster for RNA-seq data processing and initial analysis, and the Biomedical Sequencing Facility at CeMM for assistance with next-generation sequencing. Proteomics experiments were performed in the Mass Spectrometry facility of Max Perutz Labs; we thank Natascha Hartl and Weiqiang Chen for the help with sample preparation and measurements, and David Hollenstein for assistance with data analysis. All LC-MS data were acquired using the instrument pool of the Vienna Biocenter Core Facilities (VBCF). We also thank the IMBA Stem Cell Core Facility and especially Chukwuma Allison Agu, Michelle Foong-Sobis, Astrid Pentz, Ruth Fischer and Roman Stemberger for their assistance in ES cell targeting. Blastocyst injections were performed by Hans Christian Theussl from the IMBA Transgenic Core Facility. Finally, we would like to thank the team of the Animal Facility of the Max Perutz Labs Support GmbH as well as the team of the MedUni Wien Core Facility Laboratory Animal Breeding and Husbandry the for their permanent professional support.

