## Supplementary information for "Targeting the catalytic activity of HDAC1 in T cells protects against experimental autoimmune encephalomyelitis"

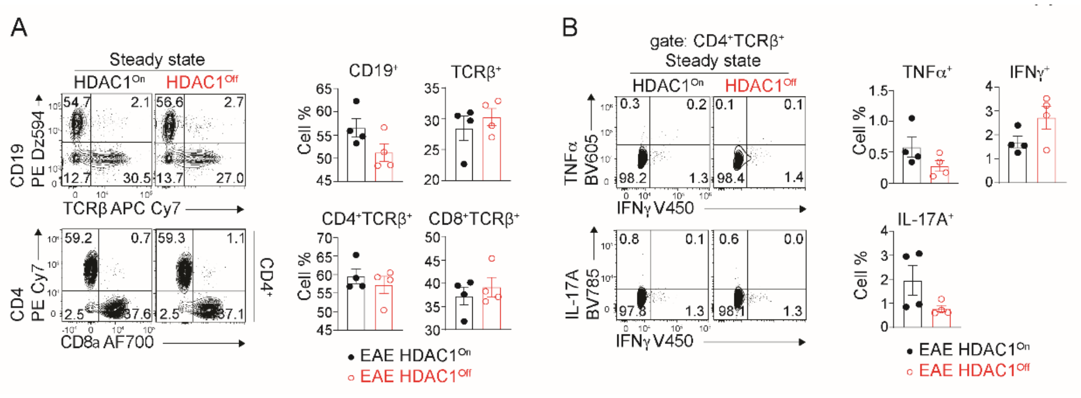

**Supplementary Figure 1. HDAC1$^{Off}$ mice show similar lymphocyte composition compared to HDAC1$^{On}$ mice at steady state. (A)** Representative flow cytometry plot showing expression of CD19, TCRβ, CD4 and CD8 in the spleen. Diagrams at the right show the summary (cell %) of all experiments. (B) Representative flow cytometry plot showing cytokine expression in splenic CD4$^{+}$ T cells upon *ex vivo* stimulation with PMA/ionomycin. The summary of all experiments is shown at the right. Data show mean ± SEM; Student t test (unpaired, two-sided with Welch's correction). Data are show representative 4 mice per genotype analyzed in 2 independent experiments.

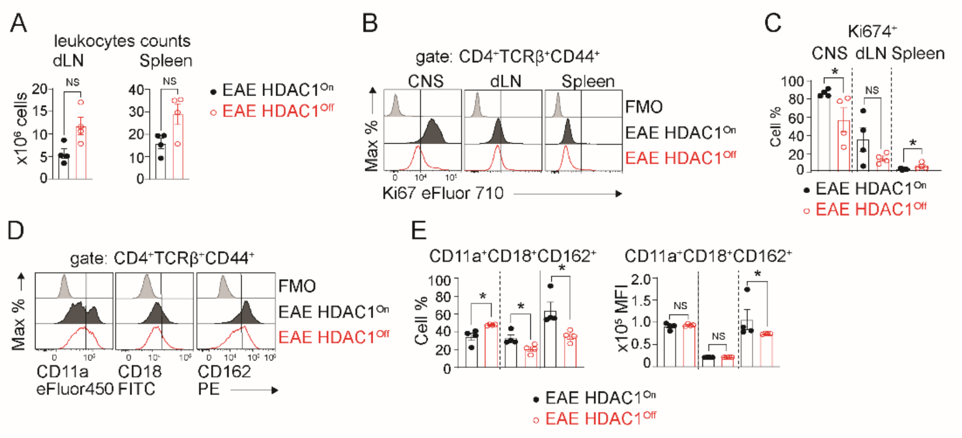

**Supplementary Figure 2. Downregulated expression of LFA-1, selectin ligands and Ki67 in HDAC1$^{Off}$ CD4$^{+}$ T cells in the EAE model.**

**(A)** Diagrams indicate the number of leukocytes present in the in the draining lymph nodes (dLN) and spleen of EAE-induced HDAC1$^{On}$ or HDAC1$^{Off}$ mice. **(B)** Representative histograms show intracellular expression of Ki67 of HDAC1$^{On}$ or HDAC1$^{Off}$ CD4$^{+}$ T cells in CNS, dLN and spleens recovered from EAE-induced mice. (**C**) Summary graph depicts frequency of CD4$^{+}$ T cells expressing Ki67 in the indicated tissue as shown in (B). **(D)** Representative histograms show cell surface expression of CD11a, CD18, CD162 on 2D2-HDAC$^{On}$ or 2D2-HDAC1$^{Off}$ CD4$^{+}$ T cells isolated from dLN and spleen recovered from EAE mice. As FMO control, CD11a, CD18 and CD162 expression on naïve CD4$^{+}$ T cells in HDAC1$^{On}$ mice is shown. (**E**) Summary graph depicts normalized geometric mean fluorescence intensity (MFI) of cell surface markers CD11a, CD18 and CD162 is shown alongside. (A,C,E) Bar diagram show mean ± SEM. *p < 0.05 (unpaired two-tailed student's t-test). NS, not significant. Data show a representative (B,D) or the summary (A,C,E) of 4 mice analyzed in 2 independent experiments.

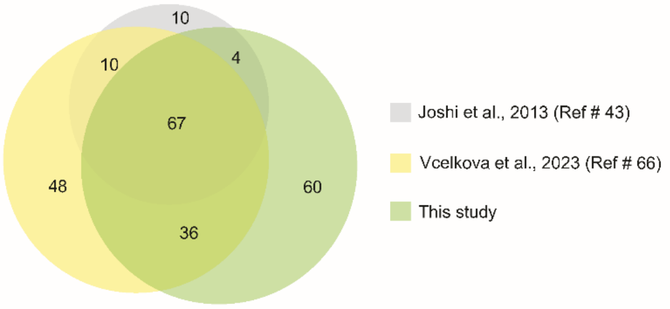

**Supplementary Figure 3. Comparison of HDAC1 interactomes**

Venn diagram illustrating the overlap between HDAC1 interactomes from three independent studies, using distinct cell types. Grey set represents HDAC1-EGFP interacting proteins identified in human CEM-T cells [1]. The yellow set represents proteins that interact with HDAC1-Flag determined in the human haploid cell line HAP1 [2]. The present study's HDAC1 interactome is indicated in green.

**Supplementary Table 1.**

List of proteins interacting with HDAC1 (I.e. HDAC1$^{On}$) in Th1 cells. The proteins are sorted according the log 2 fold change (log2FC) of enrichment compared to the *Cd4-Cre*-negative samples that were used as controls for the FLAG-IP.

| Protein IDs | Protein names | Gene names | Log2FC_HDAC1$^{On}$ / *Cd4-Cre*$^{-}$ |
|---|---|---|---|
| Q6ZQ88 | Lysine-specific histone demethylase 1A | Kdm1a | 12,84 |
| Q5UAK0 | Mesoderm induction early response protein 1 | Mier1 | 12,83 |
| Q9Z2E1 | Methyl-CpG-binding domain protein 2 | Mbd2 | 11,90 |
| Q60520 | Paired amphipathic helix protein Sin3a | Sin3a | 11,41 |
| Q8CFE3 | REST corepressor 1 | Rcor1 | 11,20 |
| Q99LB0 | Deoxynucleotidyltransferase terminal-interacting protein 1 | Dnttip1 | 11,13 |
| A2CG63 | AT-rich interactive domain-containing protein 4B | Arid4b | 10,98 |
| Q924K8 | Metastasis-associated protein MTA3 | Mta3 | 10,76 |
| F8VPQ2 | AT-rich interactive domain-containing protein 4A | Arid4a | 10,49 |
| Q8BR65 | Sin3 histone deacetylase corepressor complex component SDS3 | Suds3 | 10,28 |
| Q3U3C9 | Genetic suppressor element 1 | Gse1 | 10,14 |
| E9Q2I4 | Mitotic deacetylase-associated SANT domain protein | Elmsan1 | 10,11 |
| Q8K4B0 | Metastasis-associated protein MTA1 | Mta1 | 10,01 |
| Q6ZPK0 | PHD finger protein 21A | Phf21a | 9,86 |
| Q9R190 | Metastasis-associated protein MTA2 | Mta2 | 9,77 |
| Q9Z2D8 | Methyl-CpG-binding domain protein 3 | Mbd3 | 9,66 |
| O35207;Q9CPY4 | Cyclin-dependent kinase 2-associated protein 1;Cyclin-dependent kinase 2-associated protein 2 | Cdk2ap1;Cdk2ap2 | 9,49 |
| Q8CHY6 | Transcriptional repressor p66 alpha | Gatad2a | 9,40 |
| B1AR17 | DNA helicase | Chd3 | 9,28 |
| Q60972 | Histone-binding protein RBBP4 | Rbbp4 | 9,07 |
| Q62141 | Paired amphipathic helix protein Sin3b | Sin3b | 9,03 |
| Q8VHR5 | Transcriptional repressor p66-beta | Gatad2b | 9,00 |
| Q9CU65;Q3U2E2 | Zinc finger MYM-type protein 2 | Zmym2 | 9,00 |
| P70288 | Histone deacetylase 2 | Hdac2 | 8,98 |
| Q3U0X6;Q8K083 | Zinc finger protein 217 | Zfp217 | 8,91 |
| O09106 | Histone deacetylase 1 | Hdac1 | 8,87 |
| Q8C8M1 | SIN3-HDAC complex-associated factor | Fam60a | 8,77 |
| Q0VEE6 | Zinc finger protein 800 | Znf800 | 8,64 |
| Q9DC33 | High mobility group protein 20A | Hmg20a | 8,56 |
| Q60973 | Histone-binding protein RBBP7 | Rbbp7 | 8,38 |
| Q9Z104 | SWI/SNF-related matrix-associated actin-dependent regulator of chromatin subfamily E member 1-related | Hmg20b | 8,09 |
| Q8BM75 | AT-rich interactive domain-containing protein 5B | Arid5b | 8,08 |
| Q80TZ9 | Arginine-glutamic acid dipeptide repeats protein | Rere | 8,04 |
| Q8BYA0 | Tubulin-specific chaperone D | Tbcd | 8,02 |
| Q99N20 | Breast cancer metastasis-suppressor 1 homolog | Brms1 | 7,93 |
| Q8BIH0 | Histone deacetylase complex subunit SAP130 | Sap130 | 7,53 |

| | | | |
|---|---|---|---|
| Q6PDQ2 | Chromodomain-helicase-DNA-binding protein 4 | Chd4 | 7,51 |
| Q64321 | Zinc finger and BTB domain-containing protein 7B | Zbtb7b | 7,44 |
| Q3U1T3 | Breast cancer metastasis-suppressor 1-like protein | Brms1l | 7,39 |
| Q9EPQ8;Q3UT76 | Transcription factor 20 | Tcf20 | 7,18 |
| Q8BXJ2 | Transcriptional-regulating factor 1 | Trerf1 | 7,16 |
| Q61687 | Transcriptional regulator ATRX | Atrx | 7,10 |
| Q3U3N0 | Mesoderm induction early response protein 2 | Mier2 | 6,99 |
| Q3UHF3 | Mesoderm induction early response protein 3 | Mier3 | 6,94 |
| O88574 | Histone deacetylase complex subunit SAP30 | Sap30 | 6,89 |
| Q99PV8;Q9QX96 | B-cell lymphoma/leukemia 11B | Bcl11b | 6,88 |
| Q9WUF3 | CASP8-associated protein 2 | Casp8ap2 | 6,82 |
| Q5SX39;Q5SX40 | Myosin-4;Myosin-1 | Myh4;Myh1 | 6,81 |
| Q920S3 | GATA zinc finger domain-containing protein 1 | Gatad1 | 6,69 |
| Q62137 | Tyrosine-protein kinase JAK3 | Jak3 | 6,67 |
| Q9CR55 | UPF0547 protein C16orf87 homolog | sp\|Q9CR55\|CP087_MOUSE | 6,44 |
| Q9JLM4 | Zinc finger MYM-type protein 3 | Zmym3 | 6,30 |
| Q9R0G7 | Zinc finger E-box-binding homeobox 2 | Zeb2 | 6,30 |
| Q64318 | Zinc finger E-box-binding homeobox 1 | Zeb1 | 6,25 |
| E9Q6W4 | Zinc finger protein 296 | Zfp296 | 6,12 |
| Q3V3W4 | Zinc finger and BTB domain-containing 2 | Zbtb2 | 6,07 |
| Q6IQX8 | Zinc finger protein 219 | Zfp219 | 6,07 |
| Q5SQF8 | Histone deacetylase complex subunit SAP30L | Sap30l | 6,06 |
| Q5SPL2 | PHD finger protein 12 | Phf12 | 5,93 |
| Q497V6 | Bromo adjacent homology domain-containing 1 protein | Bahd1 | 5,88 |
| A2A484 | Zinc finger, MYND-type-containing 8 | Zmynd8 | 5,81 |
| O35344 | Importin subunit alpha-4 | Kpna3 | 5,72 |
| Q6PGA0 | REST corepressor 3 | Rcor3 | 5,58 |
| Q8BHZ4 | Zinc finger protein 592 | Znf592 | 5,55 |
| O35343 | Importin subunit alpha-3 | Kpna4 | 5,51 |
| E9Q4T4 | Coiled-coil domain-containing 71-like | Ccdc71l | 5,51 |
| P61965 | WD repeat-containing protein 5 | Wdr5 | 5,50 |
| Q3UCQ1 | Forkhead box protein K2 | Foxk2 | 5,50 |
| Q9WTK2 | Chromodomain Y-like protein | Cdyl | 5,50 |
| Q5DW34 | Histone-lysine N-methyltransferase EHMT1 | Ehmt1 | 5,48 |
| O88712 | C-terminal-binding protein 1 | Ctbp1 | 5,36 |
| A2A654 | Bromodomain PHD finger transcription factor | Bptf | 5,30 |
| P28574 | Protein max | Max | 5,30 |
| P27661;Q64522 | Histone H2AX;Histone H2A type 2-B | H2afx;Hist2h2ab | 5,28 |
| Q5DTH5 | Teashirt homolog 1 | Tshz1 | 5,24 |
| Q8C796 | REST corepressor 2 | Rcor2 | 5,18 |
| O54833 | Casein kinase II subunit alpha | Csnk2a2 | 5,17 |
| O88939 | Zinc finger and BTB domain-containing protein 7A | Zbtb7a | 4,93 |
| A2AWL7 | MAX gene-associated protein | Mga | 4,90 |

| | | | |
|---|---|---|---|
| Q8CCJ9 | PHD finger protein 20-like protein 1 | Phf20l1 | 4,88 |
| Q99KZ6 | Zinc finger protein 639 | Znf639 | 4,84 |
| Q3UH06 | Ras-responsive element-binding protein 1 | Rreb1 | 4,80 |
| Q6PAL0 | BEN domain-containing protein 3 | Bend3 | 4,79 |
| Q60960 | Importin subunit alpha-5 | Kpna1 | 4,66 |
| P50540 | Max-interacting protein 1 | Mxi1 | 4,55 |
| Q61624 | Zinc finger protein 148 | Znf148 | 4,25 |
| Q03267 | DNA-binding protein Ikaros | Ikzf1 | 4,24 |
| Q6DIC0 | Probable global transcription activator SNF2L2 | Smarca2 | 4,22 |
| Q9JMD1 | Scm-like with four MBT domains protein 1 | Sfmbt1 | 4,19 |
| Q03347 | Runt-related transcription factor 1 | Runx1 | 4,18 |
| Q6PHP4 | Zinc finger protein 512B (Fragment) | Znf512b | 4,16 |
| P42128 | Forkhead box protein K1 | Foxk1 | 4,08 |
| Q505G8 | Zinc finger protein 827 | Znf827 | 4,05 |
| O35615 | Zinc finger protein ZFPM1 | Zfpm1 | 3,99 |
| Q9Z148 | Histone-lysine N-methyltransferase EHMT2 | Ehmt2 | 3,97 |
| O08900 | Zinc finger protein Aiolos | Ikzf3 | 3,97 |
| Q3UHD9 | Arf-GAP with GTPase, ANK repeat and PH domain-containing protein 2 | Agap2 | 3,94 |
| Q9WU42 | Nuclear receptor corepressor 2 | Ncor2 | 3,84 |
| E9PYL2 | Proline-rich protein 12 | Prr12 | 3,84 |
| P50538 | Max dimerization protein 1 | Mxd1 | 3,83 |
| Q8R080 | G2 and S phase-expressed protein 1 | Gtse1 | 3,79 |
| P70121 | Zinc fingers and homeoboxes protein 1 | Zhx1 | 3,71 |
| Q64131;Q08775 | Runt-related transcription factor 3 | Runx3 | 3,67 |
| Q3U108 | AT-rich interactive domain-containing protein 5A | Arid5a | 3,65 |
| Q9ESK4 | Inhibitor of growth protein 2 | Ing2 | 3,62 |
| O08750 | Nuclear factor interleukin-3-regulated protein | Nfil3 | 3,60 |
| Q91VL9 | Zinc finger and BTB domain-containing protein 1 | Zbtb1 | 3,52 |
| E9QA22 | Zinc finger protein 644 | Zfp644 | 3,52 |
| Q3UXZ9 | Lysine-specific demethylase 5A | Kdm5a | 3,51 |
| P67871 | Casein kinase II subunit beta | Csnk2b | 3,49 |
| Q9D5D8 | Chromodomain Y-like protein 2 | Cdyl2 | 3,46 |
| Q9QZS8 | SH2 domain-containing protein 3C | Sh2d3c | 3,44 |
| Q8BZ47 | Zinc finger protein 609 | Znf609 | 3,36 |
| E9Q9M8 | PWWP domain-containing protein 2B | Pwwp2b | 3,36 |
| Q925H1 | Zinc finger transcription factor Trps1 | Trps1 | 3,31 |
| P62141 | Serine/threonine-protein phosphatase PP1-beta catalytic subunit | Ppp1cb | 3,31 |
| Q9D2D7 | Zinc finger protein 687 | Znf687 | 3,26 |
| P59178 | Lethal(3)malignant brain tumor-like protein 2 | L3mbtl2 | 3,23 |
| Q60737 | Casein kinase II subunit alpha | Csnk2a1 | 3,22 |
| A2A791 | Zinc finger MYM-type protein 4 | Zmym4 | 2,92 |
| Q6PCN7 | Helicase-like transcription factor | Hltf | 2,90 |
| Q8CGK3 | Lon protease homolog, mitochondrial | Lonp1 | 2,88 |
| Q08024 | Core-binding factor subunit beta | Cbfb | 2,88 |
| Q8VBW5 | HMG box transcription factor BBX | Bbx | 2,87 |

| | | | |
|---|---|---|---|
| Q8C208 | Zinc finger protein Eos | Ikzf4 | 2,81 |
| O35658 | Complement component 1 Q subcomponent-binding protein, mitochondrial | C1qbp | 2,80 |
| P80313 | T-complex protein 1 subunit eta | Cct7 | 2,75 |
| P80317;Q61390 | T-complex protein 1 subunit zeta | Cct6a | 2,68 |
| P80315 | T-complex protein 1 subunit delta | Cct4 | 2,67 |
| P80318 | T-complex protein 1 subunit gamma | Cct3 | 2,67 |
| P11983 | T-complex protein 1 subunit alpha | Tcp1 | 2,63 |
| O88286 | Protein Wiz | Wiz | 2,61 |
| P80316 | T-complex protein 1 subunit epsilon | Cct5 | 2,60 |
| P42932 | T-complex protein 1 subunit theta | Cct8 | 2,57 |
| Q8BTI7 | Serine/threonine-protein phosphatase 6 regulatory ankyrin repeat subunit C | Ankrd52 | 2,55 |
| P39428 | TNF receptor-associated factor 1 | Traf1 | 2,54 |
| P52293 | Importin subunit alpha-1 | Kpna2 | 2,52 |
| Q8R4E9 | DNA replication factor Cdt1 | Cdt1 | 2,51 |
| P59110 | Sentrin-specific protease 1 | Senp1 | 2,44 |
| Q8BX09 | Retinoblastoma-binding protein 5 | Rbbp5 | 2,37 |
| Q3U1Y4 | DENN domain-containing protein 4B | Dennd4b | 2,34 |
| P63038 | 60 kDa heat shock protein, mitochondrial | Hspd1 | 2,31 |
| P80314 | T-complex protein 1 subunit beta | Cct2 | 2,27 |
| Q9Z103 | Activity-dependent neuroprotector homeobox protein | Adnp | 2,18 |
| Q8CGP1;Q6ZWY9;Q64478;P10853;Q8CGP2;Q64525;Q64475;P10854;Q9D2U9;Q8CGP0;Q64524;P70696 | Histone H2B type 1-K;Histone H2B type 1-C/E/G;Histone H2B type 1-H | Hist1h2bk;Hist1h2bc;Hist1h2bh;Hist1h2bf;Hist1h2bp;Hist2h2bb;Hist1h2bb;Hist1h2bm;Hist3h2ba;Hist3h2bb;Hist2h2be;Hist1h2ba | 2,15 |
| Q7TT37 | Elongator complex protein 1 | Ikbkap | 2,11 |
| P63168;Q80ZS7 | Dynein light chain 1, cytoplasmic;Dynein light chain | Dynll1;BC048507 | 2,06 |
| Q62280 | Protein SSXT | Ss18 | 1,98 |
| Q9QXV3 | Inhibitor of growth protein 1 | Ing1 | 1,97 |
| P51174 | Long-chain specific acyl-CoA dehydrogenase, mitochondrial | Acadl | 1,95 |
| A2AJK6 | Chromodomain-helicase-DNA-binding protein 7 | Chd7 | 1,83 |
| C0HKE1;C0HKE3;C0HKE8;C0HKE7;C0HKE9;C0HKE5;C0HKE4;C0HKE2;C0HKE6;Q8CGP4;Q8R1M2;Q8BFU2;Q8CGP5;Q8CGP7;Q8CGP6;Q6GSS7;Q64523 | Histone H2A type 1-B;Histone H2A type 1-D;Histone H2A type 1-O | Hist1h2aa;H2afj;Hist3h2a;Hist1h2af;Hist1h2ak;Hist1h2ah;Hist2h2aa1;Hist2h2ac | 1,80 |
| O08664;Q921K9;Q9CXE2 | B-cell CLL/lymphoma 7 protein family member C | Bcl7c | 1,72 |

| | | | |
|---|---|---|---|
| Q99LI5 | Zinc finger protein 281 | Znf281 | 1,68 |
| P23198 | Chromobox protein homolog 3 | Cbx3 | 1,65 |
| E9Q3G8 | Nucleoporin 153 | Nup153 | 1,65 |
| Q3TKT4 | Transcription activator BRG1 | Smarca4 | 1,63 |
| Q3THW5;P0C0S6 | Histone H2A.V;Histone H2A.Z | H2afv;H2afz | 1,62 |
| Q6V595 | Kelch-like protein 6 | Klhl6 | 1,61 |
| Q9CWM4 | Prefoldin subunit 1 | Pfdn1 | 1,29 |
| Q6ZWN5 | 40S ribosomal protein S9 | Rps9 | 1,28 |
| P61759 | Prefoldin subunit 3 | Vbp1 | 1,25 |
| P62806 | Histone H4 | Hist1h4a | 1,24 |
| Q922F4 | Tubulin beta-6 chain | Tubb6 | 1,13 |
| Q9JIH2 | Nuclear pore complex protein Nup50 | Nup50 | 1,07 |
| Q8K0D0;Q04735;Q04899 | Cyclin-dependent kinase 17 | Cdk17 | 1,06 |
| Q60611;Q8VI24 | DNA-binding protein SATB1 | Satb1 | 1,03 |

**Supplementary Table 2. Differentially expressed genes**

Table showing genes differentially expressed between *in vivo* activated 2D2-HDAC1$^{Off}$ or 2D2-HDAC1$^{On}$ CD4$^{+}$ T cells. The blue and the red bars at the right indicate the Top 100 downregulated and upregulated genes in 2D2-HDAC1$^{Off}$ CD4$^{+}$ T cells, respectively, that have been used for Gene Set Enrichment Analysis (GSEA) shown in Fig. 4C.

| | gene_biotype | Pvalue HDAC1$^{Off}$ vs HDAC1$^{On}$ | padj HDAC1$^{Off}$ vs HDAC1$^{On}$ | log2FC HDAC1$^{Off}$ vs HDAC1$^{On}$ |
|---|---|---|---|---|
| Gm45552 | antisense | 9,21E-12 | 1,13E-08 | -2,802605107 |
| Saal1 | protein_coding | 3,47E-11 | 3,79E-08 | -2,788802175 |
| Gm4864 | processed_pseudogene | 1,76E-10 | 1,33E-07 | -2,770361536 |
| Gm42031 | lincRNA | 4,14E-07 | 9,68E-05 | -2,715927256 |
| Dglucy | protein_coding | 1,34E-06 | 0,00023095 | -2,514767179 |
| Acp2 | protein_coding | 1,57E-09 | 1,03E-06 | -2,431797159 |
| Ifng | protein_coding | 1,49E-07 | 4,72E-05 | -2,164153431 |
| Jarid2 | protein_coding | 1,90E-06 | 0,00029184 | -2,067115036 |
| Hdac1-ps | processed_pseudogene | 2,11E-07 | 6,27E-05 | -2,048502816 |
| Gm5824 | processed_pseudogene | 3,20E-07 | 8,27E-05 | -2,015904871 |
| Gm38014 | TEC | 2,98E-05 | 0,00243814 | -1,735391837 |
| Pitpnm2 | protein_coding | 1,60E-05 | 0,00159963 | -1,639512913 |
| Hdac1 | protein_coding | 5,00E-07 | 0,00010916 | -1,636142913 |
| Narf | protein_coding | 8,80E-11 | 7,85E-08 | -1,508408409 |
| Gzmb | protein_coding | 3,35E-08 | 1,22E-05 | -1,477875272 |
| Dnase1l3 | protein_coding | 3,55E-06 | 0,00047282 | -1,462615498 |
| Sh3gl1 | protein_coding | 5,53E-05 | 0,00396618 | -1,444382585 |
| Nfatc3 | protein_coding | 1,09E-05 | 0,00122142 | -1,440131879 |
| Cd7 | protein_coding | 1,29E-05 | 0,00137936 | -1,402454623 |
| Zfp617 | protein_coding | 2,69E-05 | 0,00227543 | -1,390004447 |
| Gm37109 | TEC | 0,00012466 | 0,00746623 | -1,310096448 |
| Anxa6 | protein_coding | 1,71E-21 | 1,68E-17 | -1,273731553 |
| Lgals3 | protein_coding | 2,62E-05 | 0,00224336 | -1,261473738 |
| Ppme1 | protein_coding | 5,88E-05 | 0,00409351 | -1,213964405 |
| Gm38115 | sense_intronic | 6,98E-05 | 0,00469191 | -1,189532278 |
| Pglyrp1 | protein_coding | 7,06E-07 | 0,00013597 | -1,170614483 |
| Dcaf12 | protein_coding | 4,44E-05 | 0,00337903 | -1,142336509 |
| Ifi213 | protein_coding | 0,00023191 | 0,01161694 | -1,137546425 |
| Gm9517 | processed_pseudogene | 5,16E-07 | 0,00011009 | -1,11755238 |
| Ms4a4b | protein_coding | 2,44E-05 | 0,00217688 | -1,104088621 |
| Tbc1d14 | protein_coding | 0,00014224 | 0,00807254 | -1,083110367 |
| Eif4g3 | protein_coding | 8,41E-05 | 0,00550736 | -1,064087211 |
| Cdk5r1 | protein_coding | 6,23E-05 | 0,00425839 | -1,059324539 |
| Slfn1 | protein_coding | 1,14E-05 | 0,00124037 | -1,058078272 |
| Tet2 | protein_coding | 0,00029428 | 0,01404079 | -1,052294384 |
| Tgfbr1 | protein_coding | 0,00032787 | 0,01528255 | -1,035452322 |

| | | | | |
|---|---|---|---|---|
| Casd1 | protein_coding | 0,00034052 | 0,01562256 | -1,031995751 |
| Urgcp | protein_coding | 0,00037187 | 0,01690304 | -1,013026235 |
| Gimap7 | protein_coding | 4,71E-05 | 0,00352954 | -0,989971017 |
| Slc25a14 | protein_coding | 2,47E-05 | 0,00218866 | -0,985497892 |
| Trib2 | protein_coding | 2,21E-05 | 0,00200538 | -0,978614669 |
| Dhx58 | protein_coding | 1,13E-05 | 0,00124037 | -0,967453212 |
| Selplg | protein_coding | 1,58E-08 | 6,47E-06 | -0,960825922 |
| Tnfrsf4 | protein_coding | 1,58E-08 | 6,47E-06 | -0,954634125 |
| Mri1 | protein_coding | 3,94E-05 | 0,00302182 | -0,94522638 |
| Utrn | protein_coding | 3,54E-06 | 0,00047282 | -0,940617163 |
| Phf11b | protein_coding | 3,57E-07 | 8,99E-05 | -0,922418335 |
| Asap1 | protein_coding | 6,41E-05 | 0,0043398 | -0,916805 |
| Setd3 | protein_coding | 0,00022232 | 0,01130975 | -0,89953803 |
| Acbd5 | protein_coding | 1,23E-06 | 0,00021542 | -0,89753794 |
| Pggt1b | protein_coding | 1,01E-06 | 0,00018401 | -0,897140095 |
| Sesn3 | protein_coding | 0,00010028 | 0,00629194 | -0,894838313 |
| Acsbg1 | protein_coding | 0,00058928 | 0,02458363 | -0,888127002 |
| Emc10 | protein_coding | 4,82E-06 | 0,000614 | -0,883131203 |
| BC051226 | bidirectional_promoter_lncRNA | 0,00061702 | 0,02517268 | -0,880765061 |
| Ecsit | protein_coding | 0,00054127 | 0,02320608 | -0,865145103 |
| Pim1 | protein_coding | 1,42E-05 | 0,00146771 | -0,861611739 |
| Clk1 | protein_coding | 1,09E-05 | 0,00122142 | -0,852707095 |
| Parp14 | protein_coding | 0,00029477 | 0,01404079 | -0,838834071 |
| S100a4 | protein_coding | 0,0006509 | 0,02597796 | -0,837257195 |
| Ifi214 | protein_coding | 7,60E-05 | 0,00503994 | -0,822062949 |
| Gm11346 | transcribed_processed_pseudoge ne | 2,04E-05 | 0,0019038 | -0,807500692 |
| Myo6 | protein_coding | 0,00033683 | 0,01552566 | -0,788096054 |
| Haus5 | protein_coding | 0,00022027 | 0,01126359 | -0,785473816 |
| Vgll4 | protein_coding | 0,00043417 | 0,01937567 | -0,783751953 |
| Adk | protein_coding | 0,0001606 | 0,00875969 | -0,77759482 |
| Terf1 | protein_coding | 0,00012748 | 0,00758525 | -0,774834424 |
| Clasp1 | protein_coding | 0,00106845 | 0,03416957 | -0,767877797 |
| Dync1li2 | protein_coding | 0,00083072 | 0,0295507 | -0,764538923 |
| Furin | protein_coding | 0,00013342 | 0,00780137 | -0,761738266 |
| Ecd | protein_coding | 0,0008134 | 0,0292219 | -0,76082117 |
| Itgb7 | protein_coding | 0,00062304 | 0,02517268 | -0,760060273 |
| Inpp5f | protein_coding | 0,00025933 | 0,01279444 | -0,754309224 |
| Tnfrsf26 | protein_coding | 6,45E-07 | 0,00012661 | -0,750996308 |
| Ankrd12 | protein_coding | 4,60E-05 | 0,00347661 | -0,748654069 |
| Rasa3 | protein_coding | 0,00075841 | 0,02778368 | -0,744538091 |
| Nabp2 | protein_coding | 0,00081552 | 0,0292219 | -0,737139824 |
| 3110056K07Ri k | processed_transcript | 0,00111216 | 0,03533721 | -0,73450259 |
| Myo1f | protein_coding | 0,00126697 | 0,0385112 | -0,732708496 |
| Pi4kb | protein_coding | 0,00080991 | 0,0292219 | -0,729354303 |
| Phf11a | protein_coding | 0,00016714 | 0,00901622 | -0,726337151 |
| Trim30a | protein_coding | 0,0001363 | 0,00787157 | -0,72625702 |
| Mrpl57 | protein_coding | 2,31E-07 | 6,49E-05 | -0,711939396 |
| Ltb | protein_coding | 6,22E-07 | 0,00012466 | -0,70803659 |

| | | | | |
|---|---|---|---|---|
| Ablim1 | protein_coding | 3,08E-09 | 1,68E-06 | -0,707971451 |
| Ifi27l2a | protein_coding | 1,07E-05 | 0,00122142 | -0,705380579 |
| Shisa5 | protein_coding | 0,00058025 | 0,02444998 | -0,702764398 |
| Ctdsp2 | protein_coding | 0,00155027 | 0,04424584 | -0,694545912 |
| Phrf1 | protein_coding | 0,00120902 | 0,03741296 | -0,689490214 |
| Supt4a | protein_coding | 2,04E-08 | 8,03E-06 | -0,685641817 |
| Ifi209 | protein_coding | 4,99E-07 | 0,00010916 | -0,683617311 |
| Atp1b3 | protein_coding | 5,25E-06 | 0,00065235 | -0,682360946 |
| Resf1 | protein_coding | 2,61E-09 | 1,51E-06 | -0,67773328 |
| Id2 | protein_coding | 0,00140255 | 0,04135206 | -0,675692726 |
| 1700012B07Rik | protein_coding | 0,00170252 | 0,04803246 | -0,673332207 |
| Snora68 | snoRNA | 0,00046188 | 0,02042686 | -0,672762627 |
| Pdcd4 | protein_coding | 3,93E-06 | 0,0005076 | -0,666012078 |
| Gm45308 | processed_pseudogene | 0,00129386 | 0,03896595 | -0,665358546 |
| Gm43305 | processed_transcript | 0,00161848 | 0,04605876 | -0,66054697 |
| Reep5 | protein_coding | 0,00092961 | 0,03147195 | -0,652957325 |
| Lsp1 | protein_coding | 1,49E-05 | 0,00150716 | -0,65080912 |
| Ms4a6b | protein_coding | 1,68E-05 | 0,00166713 | -0,647389638 |
| Phf11 | unprocessed_pseudogene | 0,00099822 | 0,0328875 | -0,640492344 |
| Lime1 | protein_coding | 0,00068689 | 0,02671279 | -0,639409876 |
| Zfp622 | protein_coding | 0,00145884 | 0,04250118 | -0,63545369 |
| Il2rg | protein_coding | 9,07E-08 | 3,07E-05 | -0,631719143 |
| Unc119b | protein_coding | 0,00149939 | 0,04313987 | -0,628155686 |
| Inpp5b | protein_coding | 0,00032844 | 0,01528255 | -0,627671041 |
| Atp8b4 | protein_coding | 0,00141494 | 0,04158741 | -0,62458749 |
| Hilpda | protein_coding | 0,00173505 | 0,04839404 | -0,618383284 |
| Smap2 | protein_coding | 0,0017934 | 0,04945945 | -0,613996046 |
| Wdr83 | protein_coding | 0,00064362 | 0,02579197 | -0,61305093 |
| Champ1 | protein_coding | 0,00093976 | 0,03161966 | -0,605682771 |
| Cd52 | protein_coding | 1,04E-07 | 3,40E-05 | -0,604966028 |
| Stk17b | protein_coding | 2,48E-06 | 0,00035836 | -0,601928342 |
| Sigirr | protein_coding | 0,00102196 | 0,03333407 | -0,595049608 |
| Ppp2r5c | protein_coding | 6,47E-06 | 0,00078368 | -0,577460042 |
| Higd2a | protein_coding | 5,53E-05 | 0,00396618 | -0,567143426 |
| Phf20l1 | protein_coding | 0,00102652 | 0,03337206 | -0,56393171 |
| Pde6d | protein_coding | 0,00027015 | 0,0131305 | -0,563023329 |
| Ddx28 | protein_coding | 0,00150273 | 0,04313987 | -0,560530073 |
| Cyb5b | protein_coding | 0,0008894 | 0,03042544 | -0,558785671 |
| Rplp0 | protein_coding | 2,13E-12 | 2,98E-09 | -0,550666599 |
| Ptpra | protein_coding | 0,00172167 | 0,0482954 | -0,550337664 |
| Eif2b2 | protein_coding | 0,00176208 | 0,04878827 | -0,538616783 |
| Gm9625 | processed_pseudogene | 1,87E-09 | 1,15E-06 | -0,538307073 |
| Cd3g | protein_coding | 2,01E-06 | 0,00030428 | -0,524308094 |
| Gimap1 | protein_coding | 1,02E-06 | 0,00018401 | -0,522487162 |
| Cd53 | protein_coding | 3,00E-07 | 7,96E-05 | -0,518815981 |
| Stk16 | protein_coding | 0,00176409 | 0,04878827 | -0,514695751 |
| Gm2a | protein_coding | 5,38E-05 | 0,00392165 | -0,513143645 |
| Gm8730 | processed_pseudogene | 1,89E-06 | 0,00029184 | -0,512834388 |

| | | | | |
|---|---|---|---|---|
| Gm6278 | processed_pseudogene | 1,38E-05 | 0,00144086 | -0,498491884 |
| Rplp0-ps1 | processed_pseudogene | 8,89E-06 | 0,00102697 | -0,493231086 |
| Gm14165 | processed_pseudogene | 2,53E-05 | 0,0022019 | -0,492683058 |
| Cd48 | protein_coding | 0,00097016 | 0,03228825 | -0,468365575 |
| Emb | protein_coding | 7,32E-09 | 3,78E-06 | -0,464446466 |
| S100a10 | protein_coding | 0,00033129 | 0,01534248 | -0,459917697 |
| Zc3hav1 | protein_coding | 0,00043967 | 0,01953248 | -0,45940792 |
| Ube2d3 | protein_coding | 2,96E-06 | 0,00042165 | -0,453711744 |
| Ube2h | protein_coding | 0,00048246 | 0,02114653 | -0,446730036 |
| Tpt1 | protein_coding | 0,00026404 | 0,01292386 | -0,441107583 |
| Eif4b | protein_coding | 0,00069059 | 0,02671279 | -0,413127821 |
| Abcg1 | protein_coding | 0,00011643 | 0,00709985 | -0,413096085 |
| Lcp2 | protein_coding | 2,83E-05 | 0,00235866 | -0,40460867 |
| Degs1 | protein_coding | 9,64E-05 | 0,00614271 | -0,393304497 |
| Vrk1 | protein_coding | 0,00013113 | 0,00775555 | -0,384381486 |
| Abracl | protein_coding | 0,00105508 | 0,03396322 | -0,377660559 |
| Cyba | protein_coding | 5,70E-07 | 0,00011657 | -0,369298309 |
| Gimap4 | protein_coding | 0,00078949 | 0,02860231 | -0,367256828 |
| Gm5869 | processed_pseudogene | 5,24E-05 | 0,00386913 | -0,366292168 |
| Rab7 | protein_coding | 2,25E-05 | 0,00202343 | -0,365302745 |
| Rpl10a | protein_coding | 0,00016638 | 0,00901622 | -0,363894135 |
| Trav9n-3 | TR_V_gene | 3,88E-05 | 0,00299912 | -0,358799973 |
| Ech1 | protein_coding | 0,00099284 | 0,03282058 | -0,358125599 |
| Traj23 | TR_J_gene | 3,22E-05 | 0,00259996 | -0,357574489 |
| Trav9n-2 | TR_V_gene | 0,000102 | 0,00633839 | -0,353623382 |
| Trav9-2 | TR_V_gene | 9,10E-05 | 0,00589881 | -0,351213187 |
| Gmfg-ps | processed_pseudogene | 0,00023456 | 0,01168966 | -0,350286382 |
| Gmfg | protein_coding | 0,00010061 | 0,00629194 | -0,345679052 |
| Eef1a1-ps1 | processed_pseudogene | 0,00032698 | 0,01528255 | -0,343856167 |
| Ppib | protein_coding | 0,00148856 | 0,04298433 | -0,340625625 |
| Saraf | protein_coding | 0,00066275 | 0,02623727 | -0,326211141 |
| Myl6 | protein_coding | 0,00091627 | 0,03112784 | -0,324250258 |
| Polr2g | protein_coding | 0,00021542 | 0,01113134 | -0,323527153 |
| Eef1a1 | protein_coding | 2,26E-06 | 0,00033581 | -0,323074362 |
| Arpc2 | protein_coding | 1,84E-06 | 0,00029184 | -0,321476247 |
| Gm13456 | processed_pseudogene | 0,00084142 | 0,02960948 | -0,319962584 |
| Elf1 | protein_coding | 0,00075066 | 0,02770661 | -0,319824693 |
| Rpl10a-ps1 | processed_pseudogene | 0,00121445 | 0,03741296 | -0,305719877 |
| Psme2 | protein_coding | 0,00020123 | 0,01045336 | -0,298975174 |
| Trac | TR_C_gene | 0,00028984 | 0,01394941 | -0,291895001 |
| Sdf4 | protein_coding | 0,00181446 | 0,04976084 | -0,288222099 |
| Rplp1 | protein_coding | 0,00053987 | 0,02320608 | -0,283280558 |
| Gm6170 | processed_pseudogene | 0,00106068 | 0,03403178 | -0,274360708 |
| Rps9 | protein_coding | 0,00014933 | 0,00828296 | -0,265448639 |
| Klf13 | protein_coding | 0,00172156 | 0,0482954 | -0,242440595 |
| Cd3d | protein_coding | 0,00116703 | 0,03657403 | -0,240154685 |
| Pigx | protein_coding | 0,0012316 | 0,03755238 | 0,283302832 |
| Cenpx | protein_coding | 0,00116971 | 0,03657403 | 0,28403215 |
| Hddc2 | protein_coding | 0,00056973 | 0,02411029 | 0,312971047 |

| | | | | |
|---|---|---|---|---|
| Uqcrb | protein_coding | 0,00060509 | 0,02488134 | 0,345575736 |
| Nudc | protein_coding | 0,00139482 | 0,04124808 | 0,362976571 |
| Cdk6 | protein_coding | 0,0014885 | 0,04298433 | 0,367982423 |
| Pdcd5 | protein_coding | 0,00013389 | 0,00780137 | 0,40097844 |
| Pdcd5-ps | processed_pseudogene | 0,00061964 | 0,02517268 | 0,403261841 |
| Nsfl1c | protein_coding | 0,00054623 | 0,0233167 | 0,405445259 |
| Gm9800 | processed_pseudogene | 0,0008636 | 0,02989192 | 0,42639257 |
| Tbcb | protein_coding | 7,17E-06 | 0,00084867 | 0,43773375 |
| Anapc5 | protein_coding | 0,00064056 | 0,02577472 | 0,439199131 |
| Cct3 | protein_coding | 9,72E-07 | 0,00018348 | 0,44235568 |
| Dynlt1a | protein_coding | 0,00129781 | 0,03896595 | 0,447123293 |
| Tspan13 | protein_coding | 0,00101934 | 0,03333407 | 0,451965271 |
| Reep3 | protein_coding | 0,00146563 | 0,04257262 | 0,464509293 |
| Gm19585 | lincRNA | 5,58E-05 | 0,00396907 | 0,464992304 |
| Eif1ax | protein_coding | 2,18E-08 | 8,23E-06 | 0,465894044 |
| Dynlt1-ps1 | processed_pseudogene | 0,00095621 | 0,03193224 | 0,466089007 |
| Tubb5 | protein_coding | 0,00035629 | 0,01626993 | 0,484588355 |
| Tuba1c | protein_coding | 0,00041101 | 0,01851034 | 0,490211923 |
| Raly | protein_coding | 0,00089572 | 0,03053545 | 0,497146942 |
| Ptma | protein_coding | 1,92E-05 | 0,00184942 | 0,5046007 |
| Ccne1 | protein_coding | 0,00017877 | 0,00948738 | 0,508192837 |
| Iltifb | protein_coding | 0,00176152 | 0,04878827 | 0,515932634 |
| Gm6682 | processed_pseudogene | 0,00070272 | 0,02684568 | 0,522903501 |
| Pdia6 | protein_coding | 2,18E-07 | 6,31E-05 | 0,531780186 |
| Mtch2 | protein_coding | 3,23E-05 | 0,00259996 | 0,540571528 |
| Tsg101 | protein_coding | 0,00072874 | 0,02751825 | 0,547001133 |
| H1f0 | protein_coding | 0,00014739 | 0,00825755 | 0,563368685 |
| Gmnn | protein_coding | 0,0008402 | 0,02960948 | 0,565723058 |
| Entpd7 | protein_coding | 0,00151888 | 0,04347619 | 0,575975342 |
| Swap70 | protein_coding | 0,00162913 | 0,04622785 | 0,576010679 |
| Slc7a11 | protein_coding | 0,00129006 | 0,03896595 | 0,581513852 |
| Gm7172 | processed_pseudogene | 0,0008592 | 0,02989192 | 0,583748231 |
| Gm15599 | processed_pseudogene | 0,00076531 | 0,02782885 | 0,593059876 |
| Comt | protein_coding | 8,08E-05 | 0,00532626 | 0,593104099 |
| Gm14308 | protein_coding | 0,0007481 | 0,02770661 | 0,59674843 |
| Tlr1 | protein_coding | 0,00109604 | 0,03493819 | 0,601458077 |
| Hmgb3-ps | processed_pseudogene | 0,00182585 | 0,04993365 | 0,602553761 |
| Trbj2-6 | TR_J_pseudogene | 0,00074434 | 0,02770661 | 0,60322584 |
| Dctpp1 | protein_coding | 0,00012472 | 0,00746623 | 0,604955013 |
| Bud13 | protein_coding | 0,001419 | 0,04158741 | 0,613621258 |
| Gm11007 | protein_coding | 0,0012156 | 0,03741296 | 0,62294614 |
| Gm11223 | processed_pseudogene | 5,01E-06 | 0,0006306 | 0,623094465 |
| Gm3756 | processed_pseudogene | 0,00025291 | 0,0125408 | 0,626549631 |
| Fdx1 | protein_coding | 0,00098369 | 0,03262808 | 0,630105098 |
| Gm14410 | protein_coding | 0,0013663 | 0,0405268 | 0,636536403 |
| Haus1 | protein_coding | 0,00075796 | 0,02778368 | 0,636880339 |
| Stmn1 | protein_coding | 2,37E-06 | 0,0003469 | 0,636978234 |
| Ubr3 | protein_coding | 0,00181195 | 0,04976084 | 0,63727234 |
| Psmd9 | protein_coding | 0,00072482 | 0,02747607 | 0,641413974 |

| | | | | |
|---|---|---|---|---|
| Rgcc | protein_coding | 9,34E-05 | 0,00599576 | 0,642665222 |
| Hmgn1 | protein_coding | 0,00014803 | 0,00825755 | 0,643241743 |
| Gm17936 | processed_pseudogene | 0,00114365 | 0,03598834 | 0,645403819 |
| Nebl | protein_coding | 0,00015782 | 0,00865622 | 0,647334428 |
| Gm14150 | processed_pseudogene | 0,00052336 | 0,02270528 | 0,647465424 |
| Tnfrsf13b | protein_coding | 3,56E-06 | 0,00047282 | 0,653815498 |
| Otulin | protein_coding | 0,00172728 | 0,04831469 | 0,654240218 |
| Tuba-rs1 | transcribed_processed_pseudogene | 0,00112911 | 0,03564499 | 0,654368685 |
| Gm3226 | processed_pseudogene | 0,00022727 | 0,01144271 | 0,66433182 |
| Gtf2a2 | protein_coding | 5,39E-05 | 0,00392165 | 0,666082373 |
| Gle1 | protein_coding | 0,00075014 | 0,02770661 | 0,666341409 |
| Ap2a2 | protein_coding | 0,001021 | 0,03333407 | 0,668093965 |
| Nup93 | protein_coding | 0,00018185 | 0,00959914 | 0,671436429 |
| Gm5620 | processed_pseudogene | 0,00026459 | 0,01292386 | 0,673034765 |
| Gm14305 | protein_coding | 0,00074927 | 0,02770661 | 0,674166259 |
| Trav14n-2 | TR_V_gene | 0,00105004 | 0,03396322 | 0,67953883 |
| Zfp52 | protein_coding | 0,00013429 | 0,00780137 | 0,680207714 |
| Mcub | protein_coding | 0,00166431 | 0,04708985 | 0,681157161 |
| Pex2 | protein_coding | 1,43E-06 | 0,00024133 | 0,688070537 |
| Tuba1b | protein_coding | 0,00015659 | 0,00863716 | 0,693865023 |
| Rcc2 | protein_coding | 0,00076163 | 0,0277979 | 0,694414314 |
| Gtpbp8 | protein_coding | 0,00132362 | 0,03949928 | 0,69729502 |
| Ptger4 | protein_coding | 0,00133173 | 0,03962108 | 0,698355607 |
| Rad51b | protein_coding | 0,00105406 | 0,03396322 | 0,702224213 |
| 9430081H08Rik | TEC | 0,00131252 | 0,03928758 | 0,702444711 |
| F730043M19Rik | bidirectional_promoter_lncRNA | 0,00122131 | 0,03747121 | 0,702453236 |
| Fuca2 | protein_coding | 0,00069912 | 0,02681234 | 0,702869189 |
| Dhfr | protein_coding | 0,00084707 | 0,02970199 | 0,704019377 |
| Ang | protein_coding | 0,00143864 | 0,04203741 | 0,70809127 |
| Nfs1 | protein_coding | 0,00086467 | 0,02989192 | 0,711547377 |
| Gm11805 | processed_pseudogene | 0,00062226 | 0,02517268 | 0,71428666 |
| Cbl | protein_coding | 0,00029603 | 0,01404079 | 0,715162908 |
| Zfp985 | protein_coding | 0,00122549 | 0,03748251 | 0,719023887 |
| Phip | protein_coding | 0,0008348 | 0,02958885 | 0,720261269 |
| Zfp599 | protein_coding | 0,00120617 | 0,03741296 | 0,722548697 |
| Cnn3 | protein_coding | 0,00032194 | 0,01519626 | 0,722689228 |
| Pacsin1 | protein_coding | 0,00060569 | 0,02488134 | 0,722743422 |
| Tspan3 | protein_coding | 0,00094041 | 0,03161966 | 0,725639765 |
| Pole3 | protein_coding | 0,00010039 | 0,00629194 | 0,726600063 |
| Ly6c2 | protein_coding | 0,00127584 | 0,03866125 | 0,726607456 |
| Gm17807 | processed_pseudogene | 0,00019477 | 0,01022597 | 0,727083565 |
| Usp40 | protein_coding | 0,00069108 | 0,02671279 | 0,728302342 |
| Dqx1 | protein_coding | 0,0008204 | 0,02928976 | 0,735323237 |
| Gnb2 | protein_coding | 0,00059093 | 0,02458363 | 0,735526235 |
| G6pc3 | protein_coding | 0,00013899 | 0,00797986 | 0,738450329 |
| Gm14434 | protein_coding | 0,00095109 | 0,0318695 | 0,738889807 |
| Vwa5a | protein_coding | 0,00119245 | 0,03716646 | 0,742157443 |

| | | | | |
|---|---|---|---|---|
| Ift46 | protein_coding | 0,00069886 | 0,02681234 | 0,745969468 |
| Azin1 | protein_coding | 1,93E-05 | 0,00184942 | 0,748676503 |
| Gm7653 | processed_pseudogene | 0,00019878 | 0,01038109 | 0,749819915 |
| E430021H15Rik | TEC | 0,00086894 | 0,02993434 | 0,751267091 |
| Zfp97 | protein_coding | 0,0011169 | 0,03537316 | 0,751551466 |
| Stag1 | protein_coding | 0,00022704 | 0,01144271 | 0,753268659 |
| Cenpc1 | protein_coding | 0,00087968 | 0,03019829 | 0,754974589 |
| Cdca4 | protein_coding | 0,00011605 | 0,00709985 | 0,756358366 |
| Slc25a24 | protein_coding | 0,00074164 | 0,02770661 | 0,766588654 |
| Por | protein_coding | 0,00059867 | 0,0248008 | 0,772147174 |
| Pus10 | protein_coding | 1,25E-05 | 0,00134869 | 0,772392229 |
| Trat1 | protein_coding | 6,09E-05 | 0,00421181 | 0,773729283 |
| Itm2a | protein_coding | 0,00012299 | 0,00745375 | 0,78571109 |
| Casp2 | protein_coding | 0,00058857 | 0,02458363 | 0,788895042 |
| Zfp39 | protein_coding | 0,00067551 | 0,02663517 | 0,789392761 |
| Lef1os1 | antisense | 0,00086366 | 0,02989192 | 0,790421898 |
| mt-Tq | Mt_tRNA | 0,00052496 | 0,02270528 | 0,794525519 |
| Gm3604 | protein_coding | 0,00068663 | 0,02671279 | 0,805973324 |
| Mtmr6 | protein_coding | 4,79E-05 | 0,00356375 | 0,811144316 |
| Usp8 | protein_coding | 7,25E-10 | 5,08E-07 | 0,814677905 |
| AU040320 | protein_coding | 0,00065374 | 0,02598534 | 0,820215329 |
| Atpif1 | protein_coding | 1,98E-05 | 0,00186795 | 0,822041938 |
| Cox7a1 | protein_coding | 1,45E-05 | 0,00148148 | 0,824495803 |
| Gm8373 | transcribed_processed_pseudogene | 0,00070907 | 0,02698318 | 0,826689369 |
| Cntrl | protein_coding | 1,87E-07 | 5,74E-05 | 0,834114251 |
| Gadd45b | protein_coding | 0,00068132 | 0,02671279 | 0,850815333 |
| Cdc20 | protein_coding | 3,67E-05 | 0,00288611 | 0,862066174 |
| Dck | protein_coding | 5,77E-05 | 0,00404842 | 0,868585909 |
| Fam122a | protein_coding | 0,00042312 | 0,01896882 | 0,87711116 |
| AI987944 | protein_coding | 0,00016812 | 0,00901954 | 0,886851702 |
| Ccdc50 | protein_coding | 7,20E-05 | 0,0048078 | 0,909338627 |
| Igfbp4 | protein_coding | 0,00055362 | 0,02353001 | 0,914159506 |
| Mus81 | protein_coding | 3,84E-05 | 0,00299605 | 0,915419556 |
| Abhd13 | protein_coding | 0,00046615 | 0,02052332 | 0,928177114 |
| Gm11914 | processed_pseudogene | 8,74E-06 | 0,0010211 | 0,928228637 |
| Ndrg2 | protein_coding | 2,41E-07 | 6,57E-05 | 0,933822543 |
| Pdcl | protein_coding | 5,69E-05 | 0,00401891 | 0,94254422 |
| Dele1 | protein_coding | 0,00048729 | 0,02126337 | 0,943117564 |
| Trav14d-3-dv8 | TR_V_gene | 3,35E-06 | 0,00046638 | 0,950102148 |
| Ercc3 | protein_coding | 0,00010893 | 0,00672622 | 0,971728051 |
| Ccr8 | protein_coding | 2,93E-05 | 0,00241846 | 0,978733271 |
| Csrnp2 | protein_coding | 0,00040034 | 0,0181131 | 0,979614318 |
| B020010K11Rik | lincRNA | 2,07E-05 | 0,0019188 | 0,981932231 |
| Klrc3 | protein_coding | 0,00017699 | 0,0094438 | 0,986334441 |
| Mysm1 | protein_coding | 1,03E-06 | 0,00018401 | 0,990508812 |
| Gm26771 | antisense | 4,77E-07 | 0,00010899 | 1,002572989 |
| Tceal9 | protein_coding | 2,80E-05 | 0,00234603 | 1,008902858 |

| | | | | |
|---|---|---|---|---|
| Trav8n-2 | TR_V_gene | 0,00027671 | 0,01338288 | 1,010388518 |
| Gnb4 | protein_coding | 1,88E-05 | 0,00184206 | 1,01808339 |
| H2-Ob | protein_coding | 6,74E-06 | 0,00080725 | 1,048922432 |
| Tmem186 | protein_coding | 0,000141 | 0,00804855 | 1,072759213 |
| Trav14d-2 | TR_V_gene | 2,52E-05 | 0,0022019 | 1,079231063 |
| Acaa2 | protein_coding | 2,17E-05 | 0,00198685 | 1,083892295 |
| Sel1l | protein_coding | 6,25E-05 | 0,00425839 | 1,096836133 |
| Aim2 | protein_coding | 9,13E-05 | 0,00589881 | 1,138984701 |
| Irf8 | protein_coding | 0,00021906 | 0,01126028 | 1,142110411 |
| Gm14419 | protein_coding | 1,55E-06 | 0,0002564 | 1,148278514 |
| Gm14295 | protein_coding | 1,38E-05 | 0,00144086 | 1,158693498 |
| Trav14d-1 | TR_V_gene | 3,87E-07 | 9,51E-05 | 1,184449377 |
| L3mbtl3 | protein_coding | 1,88E-06 | 0,00029184 | 1,189597916 |
| Cd81 | protein_coding | 1,72E-12 | 2,82E-09 | 1,215928767 |
| Gnb5 | protein_coding | 0,00014511 | 0,00818785 | 1,228285804 |
| Gm23650 | snoRNA | 1,57E-06 | 0,0002564 | 1,238869476 |
| Mpnd | protein_coding | 1,47E-08 | 6,47E-06 | 1,246759599 |
| Pigyl | protein_coding | 4,14E-07 | 9,68E-05 | 1,255853549 |
| Gm42732 | TEC | 3,37E-06 | 0,00046638 | 1,273339901 |
| Helq | protein_coding | 2,63E-05 | 0,00224336 | 1,311903047 |
| Atad5 | protein_coding | 3,62E-05 | 0,00286261 | 1,312497785 |
| Gm14718 | antisense | 5,44E-07 | 0,00011354 | 1,358205258 |
| Lyrm7 | protein_coding | 1,94E-05 | 0,00184942 | 1,488987316 |
| Trav14n-1 | TR_V_gene | 1,31E-10 | 1,07E-07 | 1,50113695 |
| Gm37716 | TEC | 8,10E-09 | 3,98E-06 | 1,633843486 |
| Net1 | protein_coding | 3,32E-05 | 0,00264735 | 1,663883848 |
| Aaas | protein_coding | 3,84E-06 | 0,00050224 | 1,894202894 |
| Atp6v1g3 | protein_coding | 5,35E-06 | 0,00065601 | 1,909737264 |
| Enpp1 | protein_coding | 6,32E-14 | 1,55E-10 | 2,017310285 |
| Dapl1 | protein_coding | 7,72E-16 | 3,79E-12 | 2,018908999 |
| Susd1 | protein_coding | 5,26E-13 | 1,03E-09 | 2,04744085 |
| Cd72 | protein_coding | 8,32E-11 | 7,85E-08 | 2,181315854 |
| Bsdc1 | protein_coding | 1,80E-14 | 5,91E-11 | 2,229890051 |
| Ptgr1 | protein_coding | 5,46E-08 | 1,91E-05 | 2,372025144 |
| Zbtb20 | protein_coding | 9,16E-09 | 4,28E-06 | 2,421192591 |

**Supplementary Table 2: GO_BP terms**

Table showing the results of Gene Set Enrichment Analysis (GSEA) for gene sets from the Gene Ontology database term biological processes (GO:BP). A negative normalized enrichment score (NES) indicates a downregulation in $HDAC1^{Off}$ $CD4^{+}$ T cells.

| ID | Description | NES | p.adjust |
|---|---|---|---|
| GO:0034097 | response to cytokine | -1,6428703 | 0,0088757 |
| GO:0045087 | innate immune response | -1,6595766 | 0,0088757 |
| GO:0031347 | regulation of defense response | -1,6757479 | 0,0088757 |
| GO:0071345 | cellular response to cytokine stimulus | -1,6058021 | 0,0088757 |
| GO:0098542 | defense response to other organism | -1,5648023 | 0,01126871 |
| GO:0050776 | regulation of immune response | -1,5561705 | 0,02926367 |
| GO:0001816 | cytokine production | -1,5529952 | 0,03096929 |
| GO:0032101 | regulation of response to external stimulus | -1,5533246 | 0,03096929 |
| GO:0042110 | T cell activation | -1,5846591 | 0,03096929 |
| GO:0006909 | phagocytosis | -1,8126068 | 0,03096929 |
| GO:0002684 | positive regulation of immune system process | -1,4963188 | 0,03096929 |
| GO:0002831 | regulation of response to biotic stimulus | -1,6830893 | 0,03096929 |
| GO:0030335 | positive regulation of cell migration | -1,6695972 | 0,0321972 |
| GO:0003012 | muscle system process | -1,7255461 | 0,0321972 |
| GO:0060627 | regulation of vesicle-mediated transport | -1,6096771 | 0,0321972 |
| GO:0007155 | cell adhesion | -1,4828467 | 0,0321972 |
| GO:0050764 | regulation of phagocytosis | -1,9468036 | 0,03271442 |
| GO:0040017 | positive regulation of locomotion | -1,6188561 | 0,03470607 |
| GO:2000147 | positive regulation of cell motility | -1,655553 | 0,03547808 |
| GO:0001817 | regulation of cytokine production | -1,5278334 | 0,03547808 |
| GO:0003008 | system process | -1,4571487 | 0,0370228 |
| GO:0022610 | biological adhesion | -1,4805587 | 0,03758623 |
| GO:0001818 | negative regulation of cytokine production | -1,6925764 | 0,04382252 |
| GO:0000278 | mitotic cell cycle | 1,40349717 | 0,04512664 |

" – " indicates downregulation in $HDAC1^{Off}$ $CD4^{+}$ T cells

**Supplementary Table 3. Enrichment of top 100 ranked genes**

Table showing the enrichment score for the Top 100 downregulated and upregulated genes in 2D2-HDAC1$^{Off}$ CD4$^{+}$ T cells in WT and 2D2-HDAC1$^{cKO}$ CD4$^{+}$ T cells as described in Fig. 4C.

| ID | setSize | NES | p.adjust | leading_edge | core_enrichment |
|---|---|---|---|---|---|
| top_down-regulated_in_HDAC1$^{Off}$ | 100 | -2,154518488 | 6,25E-10 | tags=40%, list=9%, signal=37% | Gimap7/Terf1/Foxk1/Zfp617/Clk1/Nabp2/Clasp1/Pim1/1700012B07Rik/Dync1li2/Tet2/Selplg/3110056K07Rik/Cd7/Furin/Tnfrsf26/Myo1f/Utrn/Vgll4/Itgb7/Ms4a6b/Phf11a/Ifi214/Lsp1/S100a4/Slfn1/Tnfrsf4/Ifi209/Rasa3/Tbc1d14/Phf11b/Trib2/Resf1/Pglyrp1/Ablim1/Gm45552/Id2/Ms4a4b/Hdac1/Gm4864 |
| top_up-regulated_in_HDAC1$^{Off}$ | 100 | 1,974753021 | 4,23E-06 | tags=25%, list=10%, signal=23% | Cd81/Dapl1/Gnb5/Itm2a/Pigyl/Cnn3/Gnb4/Tspan3/G6pc3/Acaa2/Susd1/Atpif1/Mpnd/Ndrg2/Pole3/Ptgr1/Cox7a1/Aaas/Ly6c2/Fuca2/Vwa5a/Tuba1b/Enpp1/Cd72/Igfbp4 |
